# Photoswitchable Molecular Glues Enable Optical Control of Transcription Factor Degradation

**DOI:** 10.1101/2023.04.09.536172

**Authors:** Christopher J. Arp, Martin Reynders, Vedagopuram Sreekanth, Praveen Kokkonda, Michele Pagano, Amit Choudhary, Dirk Trauner

## Abstract

Immunomodulatory drugs (IMiDs), which include thalidomide and its derivatives, have emerged as the standard of care against multiple myeloma. They function as molecular glues that bind to the E3 ligase cereblon (CRBN) and induce protein interactions with neosubstrates, including the transcription factors Ikaros (IKZF1) and Aiolos (IKZF3). The subsequent ubiquitylation and degradation of these transcription factors underlies the antiproliferative activity of IMiDs. Here, we introduce photoswitchable immunomodulatory drugs (PHOIMiDs) that can be used to degrade Ikaros and Aiolos in a light-dependent fashion. Our lead compound shows minimal activity in the dark and becomes an active degrader upon irradiation with violet light. It shows high selectivity over other transcription factors, regardless of its state, and could therefore be used to control the levels of Ikaros and Aiolos with high spatiotemporal precision.

---

Molecular glues were serendipitously discovered as small molecules that bring together proteins to establish or enhance a protein-protein interaction (PPI).^[1,2]^ The thalidomide derivatives lenalidomide and pomalidomide (“IMiDs”) are important treatments for acute leukemia, multiple myeloma, myelodysplastic syndromes, and Kaposi sarcoma. Long after gaining clinical significance, IMiDs were discovered to function as molecular glues. In 2010, Handa and coworkers made the seminal discovery that the teratogenicity of thalidomide is mediated by cereblon (CRBN).^[3]^ Later, it was shown that lenalidomide binds to CRBN to promote the degradation of the zinc-finger (ZF) transcription factors Ikaros (IKZF1) and Aiolos (IKZF3), explaining the antiproliferative effects of IMiDs. (Figure 1A).^[4–8]^ Furthermore, the same mechanism was shown to induce SALL4 degradation, which was discovered to mediate teratogenicity.^[9,10]^ Together, these discoveries provide clinical validation for targeted protein degradation as a therapeutic modality and have sparked the development of new, more potent IMiDs, including iberdomide (CC-220) and mezigdomide (CC-92480) (Figure 1B).^[11–13]^

**Figure 1.**
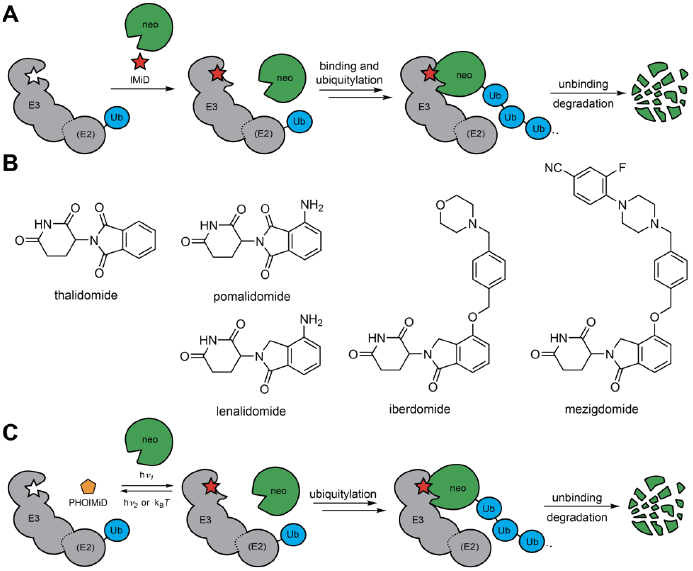
IMiDs and PHOIMiDs. (A) Schematic representation of IMiD function. The IMiD binds to an E3 ligase, inducing a conformational change in the protein structure. This change directs the recruitment of a neosubstrate (neo), which binds and forms a ternary complex with the E3 ligase and IMiD. Subsequent ubiquitylation leads to degradation of the neosubstrate. (B) Examples of established IMiDs. (C) Schematic representation of PHOIMiD function. The PHOIMiD is activated with light by incorporation of a photoswitch that allows for reversible control between the inactive form (yellow pentagon) and the active form (red star).

While IMiDs have demonstrated clinical efficacy, systemic exposure has led to adverse events such as hematologic side effects, peripheral neuropathy, and central neurotoxicity.^[14,15]^ We were inspired to make photoswitchable IMiDs that could potentially find use as precision drugs by restricting their activity to defined areas at set times with light (Figure 1C). We and others have previously developed strategies to control the activity of CRBN-based bifunctional degraders (*i*.*e*., PROTACs) with light.^[16–20]^ Building out from our previous work, we sought to make photoswitchable IMiDs that could serve as low-molecular weight degraders.

We designed photoswitchable molecular glues based on the incorporation of a minimal photoswitch into the scaffold of validated molecular glue compounds. The ideal PHOIMiDs would be inactive in the dark and remain unbound to their protein targets until activated with light to initiate binding and downstream function (Figure 1C). To date, most IMiDs are structurally related derivatives of thalidomide, lenalidomide, or pomalidomide, which feature an invariant glutarimide moiety (Figure 1B). Therefore, we started with two scaffolds, lenalidomide and pomalidomide, and extended their aromatic ring to an azobenzene in position 4 (Figure 2A). Azobenzene photoswitches are desirable owing to their large and predictable geometrical changes, their resistance to fatigue with repeated photocycling, and their tunable photophysical properties.^[21,22]^ To explore the effects of azobenzene position we also designed photoswitches where the azobenzene branched from position 5 of pomalidomide (Figure 2A, right). Potent IMiDs such as CC-885 and eragidomide (CC-90009) are lenalidomide derivatives featuring large substituents at position 5.^[11]^ Since the phthalimide aryl system contains an aniline that can be functionalized using standard approaches, these three scaffolds were well suited for “azoextension” and required only a small increase in molecular weight. Furthermore, a two-step synthesis from the corresponding phthalimide provided synthetic access to a wide range of substituted azobenzenes from diverse anilines (Figure 2B).

**Figure 2.**
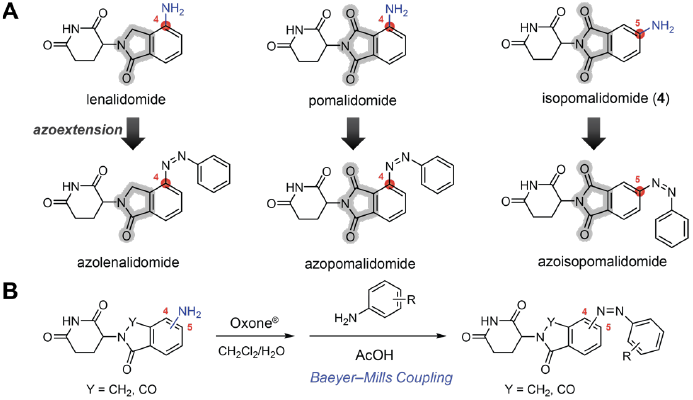
Design and synthesis of PHOIMiDs. (A) Lenalidomide and pomalidomide were designed to be modified with an azobenzene photoswitch emerging from the C4 or C5 positions of the phthalimide ring. (B) Synthesis of photoswitchable IMiDs

Satisfying key criteria for photoswitch design, we synthesized a small library of azobenzene-based PHOIMiDs (Figure 3A). Among these candidates, **PHOIMiD-9** emerged as our lead compound. The synthesis of **PHOIMiD-9** started from 4-aminophthalimide (**1**). Activation with ethyl chloroformate yielded intermediate **2** and substitution with 3-aminopiperidine-2,6-dione (**3**) yielded isopomalidomide (**4**). Intermediate **4** was oxidized with Oxone to the corresponding nitroso compound (**5**), which was used directly in a Baeyer–Mills coupling with 4-(2-methoxyphenoxy)aniline (**6**) to afford **PHOIMiD-9**. Other compounds were synthesized analogously from **4** or from lenalidomide (see Supporting Information).

**Figure 3.**
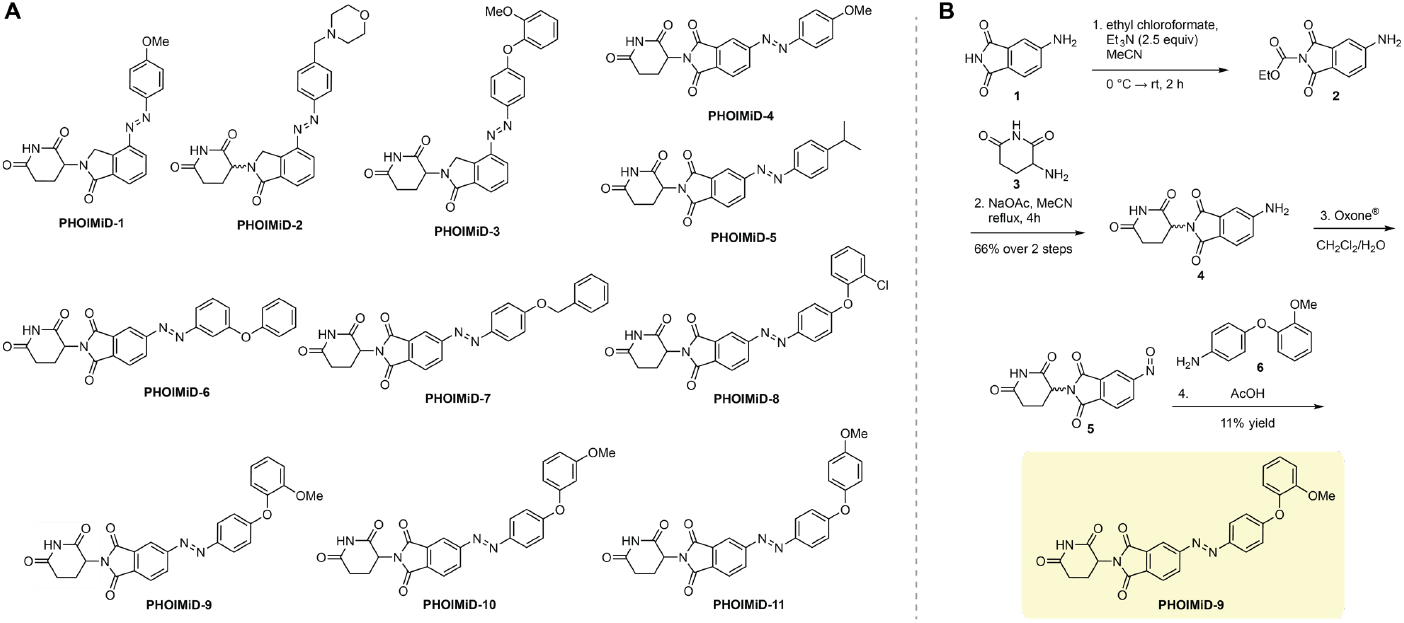
Structures and synthesis of PHOIMiDs. (A) Small library of synthesized PHOIMiDs based on lenalidomide or isopomalidomide scaffolds bearing azobenzene substitutions ranging in position, size, and hydrophobicity. (B) Synthesis of isopomalidomide (**4**) and **PHOIMiD-9**.

The photophysical properties of the lead compound, **PHOIMiD-9**, are shown in Figure 4. Photoswitching of **PHOIMiD-9** to the (*Z*) isomer was achieved by irradiation with an optimal wavelength of 390 nm (Figure 4A). (*Z*)-**PHOIMiD-9** was rapidly photoisomerized to the (*E*) isomer using irradiation with wavelengths >450 nm (Figure 4B). Under dark conditions, the metastable (*Z*) isomer of **PHOIMiD-9** relaxes back to the (*E*) isomer with a half-life of 1 h at 37 °C in DMSO (Figure 4C). Multiple cycles of photoswitching between (*Z*)-**PHOIMiD-9** and (*E*)-**PHOIMiD-9** using 390 nm and 500 nm irradiation, respectively, was accomplished without evidence of photoswitch fatigue (Figure 4D). Other PHOIMiDs showed similar photophysical properties (Figures S1-S3). Taken together, these data underscore the potential utility of **PHOIMiD-9** in experiments using low-dose, pulsed irradiation.

**Figure 4.**
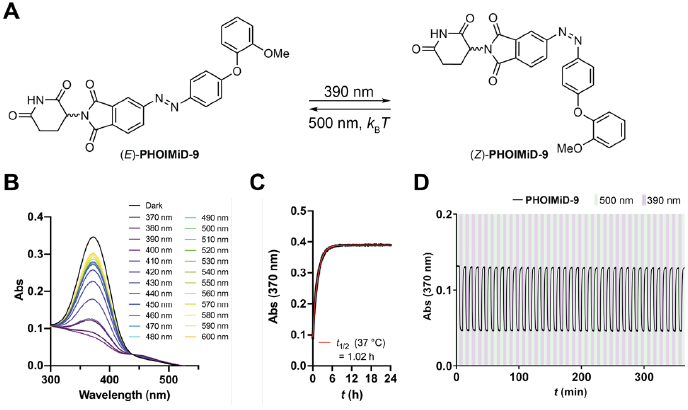
Photophysical properties of **PHOIMiD-9**. (A) **PHOIMiD-9** is photoisomerized to the (*Z*) isomer using 390 nm irradiation and isomerized to the (*E*) isomer through either thermal relaxation (*k*_B_*T*) or irradiation with 500 nm light. (B) UV-Vis absorbance scans of **PHOIMiD-9** under dark conditions or under irradiation from 370 nm to 600 nm. (C) Thermal relaxation of (*Z*)-**PHOIMiD-9** (25 μM) in DMSO at 37 °C. (D) Reversible photoswitching of **PHOIMiD-9** between its (*E*) and (*Z*) isomers upon irradiation with 500 nm or 390 nm light.

We screened PHOIMiDs for antiproliferative effects in the acute lymphoblastic leukemia cell line RS4;11. In these MTS-based cell proliferation assays, RS4;11 cells were treated with PHOIMiDs under dark or pulsed light conditions (Figure 5A, Figure S4; pulse protocol shown in Figure 6B). We tested isopomalidomide (**4**) under the same conditions and found that it did not affect cell proliferation (Figure S4L). Of the lenalidomide-derived compounds, **PHOIMiD-3** showed significant antiproliferative activity at 100 μM (<10% cell viability), but its effects were neither dose- nor light-dependent (Figure S4C). Similar results were obtained with **PHOIMiD-5, PHOIMiD-7**, and **PHOIMiD-11** (Figure S4E, S4G, S4K). In these assays, **PHOIMiD-10** showed low cytotoxicity at the highest doses (about 40% cell viability at 100 μM; Figure S4J), whereas **PHOIMiD-1, PHOIMiD-2**, and **PHOIMiD-4** showed no effects on cell proliferation (Figure S4A, S4B, S4D). **PHOIMiD-8**, with an ortho-chloro substituted phenoxy group, displayed light-dependent inhibition of cell proliferation with an IC_50_ of 11 μM, indicating that this substitution pattern was a key determinant of activity (Figure S4H). Indeed, **PHOIMiD-9**, with the chloro substituent replaced by a methoxy group, displayed an IC_50_ of 4 μM and the largest window between light and dark activity (Figure 5A; Figure S4I). These features prompted us to select **PHOIMiD-9** as the lead compound for further investigations.

**Figure 5.**
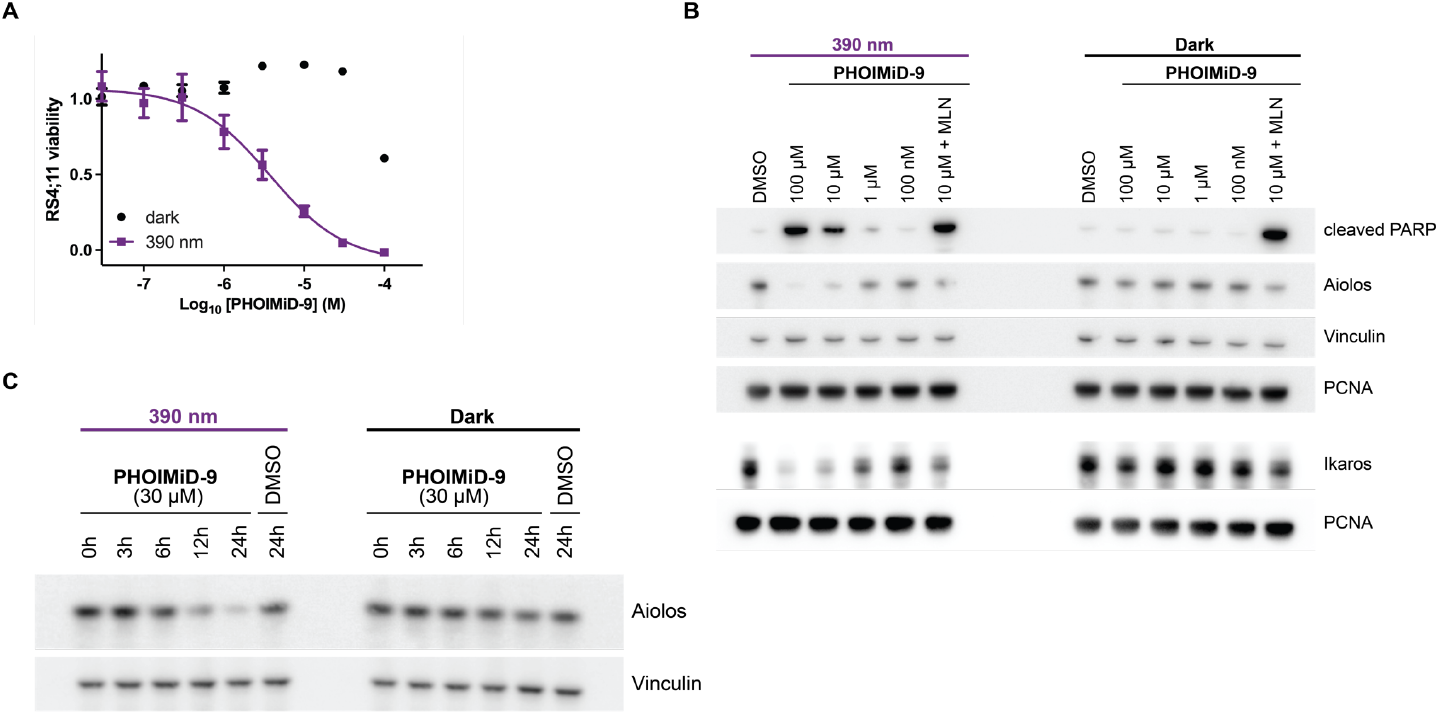
(A) Dose response curves of **PHOIMiD-9** in MTS-based cell proliferation assays. Briefly, RS4;11 cells were treated with DMSO (1%) or increasing concentrations of compound under dark or pulsed irradiation conditions for 72 hours and cell viability was assessed using MTS assays. Data are the mean ± SEM (*n* = 3). (B) Immunoblots from lysates of RS4;11 cells treated with **PHOIMiD-9** for 24 hours under dark or pulsed irradiation conditions in a dosing range from 100 nM to 100 μM. MLN (1 μM MLN4924) was used as an additional control. (C) Immunoblots from lysates of RS4;11 cells treated with **PHOIMiD-9** (30 μM) for 0 to 24 hours under dark or pulsed irradiation conditions.

**Figure 6.**
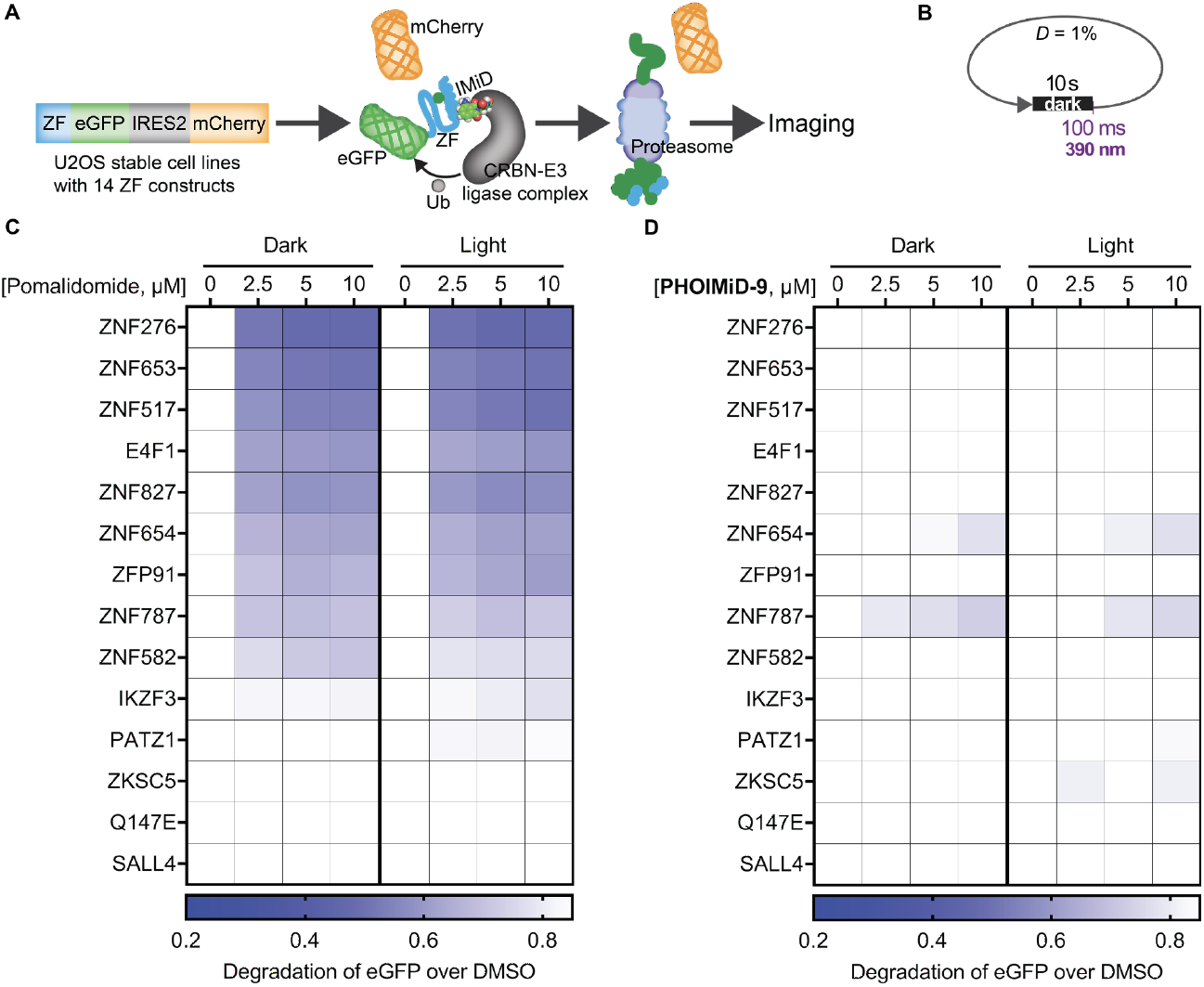
Evaluation of off-target ZF degradation of **PHOIMiD-9**. (A) Schematic of the automated imaging screen for the degradation of ZF degrons. U2OS cells stably expressing a library of 14 ZF degrons fused to eGFP enable ratiometric imaging to measure ZF degradation. (B) Schematic representation of the irradiation pulse protocol. Cells are kept under dark conditions (dark) or irradiated with brief light pulses (100 ms) following extended periods under dark conditions (10 s) for a 1% duty cycle (*D*) of irradiation. (C-D) Degradation of ZF degrons by (C) pomalidomide and (D) **PHOIMiD-9** in a dose range from 0 to 10 μM

We then probed for degradation of the main targets of IMiDs, Ikaros and Aiolos, by Western blot. RS4;11 cells were treated with DMSO or **PHOIMiD-9** in a dosing range from 100 nM to 100 μM for 24 hours under dark or pulsed light conditions (Figure 5B; pulse protocol shown in Figure 6B). We observed a substantial decrease in Ikaros and Aiolos levels upon treatment with **PHOIMiD-9** under 390 nm pulsed irradiation and a concurrent increase in cleaved PARP levels, particularly at doses between 1 μM and 100 μM, but not in the dark (Figure 5B). Similarly, in the multiple myeloma cell line U266, **PHOIMiD-9** displayed light-dependent degradation of Ikaros and Aiolos (Figure S5). Treatment with **PHOIMiD-9** (10 μM) together with the neddylation inhibitor MLN4924 (1 μM), which inhibits the activity of the cullin– RING ubiquitin ligase (CRL) complex CRL4^CRBN^, blocked the full degradation of Ikaros and Aiolos. The time dependence of Aiolos degradation in RS4;11 cells is shown in Figure 5C. Aiolos levels were substantially reduced by 12 hours of compound treatment and pulsed irradiation, and further decreased by 24 hours of compound treatment; in the dark, **PHOIMiD-9** had no effect on Aiolos levels. We also tested compound **4** under the same dark or pulsed light conditions and found that no substantial decrease in the levels of Ikaros was observed under either 390 nm or dark conditions (Figure S6).

To evaluate the potential of **PHOIMiD-9** as a selective tool compound, we performed a high-throughput screening assay to measure off-target effects of **PHOIMiD-9** in comparison to pomalidomide against a panel of validated IMiD targets (Figure 6).^[23]^ For these screens, we used U2OS cell lines stably expressing a library of validated degrons from 14 ZF targets.^[24]^ The constructs used for expression contain a ZF degron-fused eGFP in addition to a constitutively expressed mCherry reporter.

Using this system, ZF degradation can be measured by ratiometric imaging of eGFP versus mCherry signal (Figure 6A). Briefly, cells were incubated under dark conditions, or with pulsed illumination of 390 nm (100 ms per 10 s, 1% duty cycle) for 24 hours (Figure 6B) upon treatment with pomalidomide (Figure 6C) or **PHOIMiD-9** (Figure 6D). Pomalidomide induced substantial degradation of most ZF off-targets and the extent of ZF degradation did not differ between dark and light conditions (Figure 6C). On the other hand, **PHOIMiD-9** led to very slight degradation of only a few of the off-targets we probed, with subtle differences in the extent of degradation between dark and light conditions (Figure 6D). Recently, we demonstrated reduced off-targets with 5-substituted degraders,^[24]^ and another recent study shows that blocking the 5-position of the phthalimide in the IMiD scaffold prevents the formation of toxic 5-OH metabolites.^[25]^ Taken together, these data suggest that the 5-substituted phthalimide of **PHOIMiD-9** reduces off-target degradation relative to pomalidomide. These results are consistent with structure-activity relationships from other studies that have found reduced off-targets profiles associated with 5-substituted degraders.

In summary, we have described the design, synthesis, and biological evaluation of photoswitchable IMiDs, termed **PHOIMiDs**. The 5-substituted pomalidomide derivative **PHOIMiD-9** emerged as a lead compound. **PHOIMiD-9** is activated by 390 nm light while remaining inactive in the dark. Light-dependent inhibition of cell proliferation in RS4;11 cells was demonstrated using **PHOIMiD-9** with low-dose, pulsed irradiation with 390 nm light. Finally, **PHOIMiD-9** was used to optically control the degradation of the neosubstrates Ikaros and Aiolos, showing selectivity over other ZF transcription factors. Light-activatable IMiDs, such as **PHOIMiD-9**, could serve as useful tools for studying the cellular effects of transcription factor degradation with spatial and temporal precision. Additionally, these tool compounds could be used in engineered systems to optically control the degradation of proteins using IMiD-inducible degrons or to activate split protein systems employing IMiD-inducible dimerization.^[26–29]^

## Supporting Information

The authors have cited additional references within the Supporting Information.^[30]^

## Acknowledgements

This work was supported by NIH (R01 GM137606 and R01 GM132825) to A.C.

