## Supporting Information for "Photoswitchable Molecular Glues Enable Optical Control of Transcription Factor Degradation"

**Abstract:** Immunomodulatory drugs (IMiDs), which include thalidomide and its derivatives, have emerged as the standard of care against multiple myeloma. They function as molecular glues that bind to the E3 ligase cereblon (CRBN) and induce protein interactions with neosubstrates, including the transcription factors Ikaros (IKZF1) and Aiolos (IKZF3). The subsequent ubiquitylation and degradation of these transcription factors underlies the antiproliferative activity of IMiDs. Here, we introduce photoswitchable immunomodulatory drugs (PHOIMiDs) that can be used to degrade Ikaros and Aiolos in a light-dependent fashion. Our lead compound shows minimal activity in the dark and becomes an active degrader upon irradiation with violet light. It shows high selectivity over other transcription factors, regardless of its state, and could therefore be used to control the levels of Ikaros and Aiolos with high spatiotemporal precision.

#### Table of Contents

I. General Experimental ---------------------------------------------------------------------------------------- 4

II. Synthetic Procedures and Characterization ------------------------------------------------------------ 5

III. UV-Vis Spectroscopy ---------------------------------------------------------------------------------------19

Table S1. UV-Vis wavelength scan steps.----------------------------------------------------------20

IV. Cell Lines and Reagents------------------------------------------------------------------------------------21

V. Automated Imaging Screening and Analysis-----------------------------------------------------------21

Table S2. Amino acid sequences of the zinc finger motifs. -------------------------------------22

VI. Cell Proliferation Assays-----------------------------------------------------------------------------------23

VII. Immunoblotting---------------------------------------------------------------------------------------------23

Table S3. Antibody information and concentrations.---------------------------------------------25

VIII. Supplemental Figures ------------------------------------------------------------------------------------26

Figure S1. UV-Vis wavelength scans for **PHOIMiD-1–11**.----------------------------------------27

Figure S2. Thermal relaxation of PHOIMiDs in DMSO at 37 °C. -----------------------------28

Figure S3. Thermal relaxation of PHOIMiDs in 50% DMSO/PBS at 37 °C.-----------------29

Figure S4. MTS data for **PHOIMiD-1–11** and isopomalidomide (**4**).--------------------------30

Figure S5. Western blot of Ikaros and Aiolos in U266 cells treated with **PHOIMiD-9**.----31

Figure S6. Western blot of Ikaros in RS4;11 cells treated with **4**. ----------------------------32

Figure S7. Western blot of Ikaros in RS4;11 cells treated with **PHOIMiD-1**.----------------33

Figure S8. Western blot of RS4;11 cells treated with **PHOIMiD-4** or lenalidomide.-------34

Figure S9. Uncropped Western blots for Figure 5B, top. ---------------------------------------35

Figure S10. Uncropped Western blots for Figure 5B, bottom.----------------------------------36

Figure S11. Uncropped Western blots for Figure 5C. -------------------------------------------37

Figure S12. Uncropped Western blots for Figure S5.--------------------------------------------38

Figure S13. Uncropped Western blots for Figure S6.--------------------------------------------39

Figure S14. Uncropped Western blots for Figure S7.--------------------------------------------40

Figure S15. Uncropped Western blots for Figure S8.--------------------------------------------41

IX. References------------------------------------------------------------------------------------------------42

X. NMR Spectra ---------------------------------------------------------------------------------------------43

SUPPORTING INFORMATION

#### I. General Experimental

Anhydrous dichloromethane ($CH_2Cl_2$), diethyl ether ($Et_2O$), 1,4-dioxane, tetrahydrofuran (THF), toluene (PhMe), and *N*,*N*-dimethylformamide (DMF) were obtained by passing the solvent through activated alumina columns into flame-dried glassware. Other solvents and reagents were used as obtained from commercial vendors (Acros Organics, AK Scientific, Alfa Aesar, Chem-Impex International, Combi-Blocks, Sigma-Aldrich, Strem Chemicals, Synthonix, Tokyo Chemical Industry Co.) unless otherwise described. Thin-layer chromatography (TLC) was performed for reaction monitoring using silica gel 60 glass plates pre-coated with $F_{254}$ fluorescent indicator (Millipore Sigma) and visualized by blocking of ultraviolet light ($\lambda$ = 254 nm) or by staining with aqueous potassium permanganate ($KMnO_4$) solution, aqueous acidic ceric ammonium molybdate (IV) (CAM) solution, acidic ethanolic *p*-anisaldehyde solution, or butanolic ninhydrin solution, followed by gentle heating with a heat gun. Flash-column chromatography was performed at room temperature under pressure of nitrogen with silica gel (60 Å, 40-63 µm, Silicycle or Merck) using glass columns or a Teledyne Isco MPLC Combi*Flash*® Rf+. High-performance liquid chromatography (HPLC) purification was performed on an Agilent 1260 Infinity II LC with a reverse-phase (RP) Phenomenex Semipreparative Column (00D-4439-E0 Gemini, C18 phase, 3 µm particle size, 110 Å pore size) with a flow rate of 8 mL/min and solvent mixtures of 0.1% formic acid (FA) in acetonitrile (HPLC grade) and water (HPLC grade). Nuclear magnetic resonance (NMR) spectra were recorded at 25 °C in 5 mm borosilicate glass tubes (Wilmad-LabGlass). NMR samples were prepared in deuterated solvents purchased from Cambridge Isotope Laboratories (dimethyl sulfoxide-$d_6$ or DMSO-$d_6$, 99.9% D). Proton nuclear magnetic resonance ($^1$H NMR) spectra were recorded on a Bruker Avance III HD 400 MHz spectrometer equipped with a CryoProbe™ at 25 °C, are reported in parts per million (ppm, δ scale) downfield from tetramethylsilane (TMS, δ = 0 ppm), and are referenced internally to the residual protium resonances of the NMR solvent (DMSO-$d_5$ in DMSO-$d_6$, δ = 2.50 ppm). Proton-decoupled carbon-13 nuclear magnetic resonance ($^{13}$C{$^1$H} NMR) spectra were recorded on a Bruker Avance III HD 400 MHz spectrometer equipped with a CryoProbe™ at 25 °C, are reported in parts per million (ppm, δ scale) downfield from tetramethylsilane (TMS, δ = 0 ppm), and are referenced internally to the central line of carbon-13 resonances of the NMR solvent (DMSO-$d_5$ in DMSO-$d_6$, δ = 39.52 ppm). The reported data are represented as: chemical shift in parts per million (ppm, δ scale) (multiplicity, coupling constants *J* in Hz, integration). Multiplicities are abbreviated as: s, singlet; d, doublet; t, triplet; q, quartet; quint, quintet; sext, sextet; hept, heptet; br, broad; m, multiplet; or combinations thereof. Signals that cannot be clearly identified due to second-order effects, overlapping signals, or for other reasons are labeled as multiplet or broad. High-resolution mass spectrometry (HRMS) was conducted using an Agilent 6224 Accurate-Mass time-of-flight (TOF) liquid-chromatography mass spectrometer (LC/MS) in combination with either atmospheric pressure chemical ionization (APCI) or electrospray ionization (ESI) methods. Liquid chromatography-mass spectrometry (LCMS) was conducted using an Agilent Technologies 1260 II Infinity liquid chromatography system coupled to an Agilent Technologies 6120 Quadrupole mass spectrometer with an APCI ionization source.

#### II. Synthetic Procedures and Characterization

##### 5-amino-2-(2,6-dioxopiperidin-3-yl)isoindoline-1,3-dione (4)

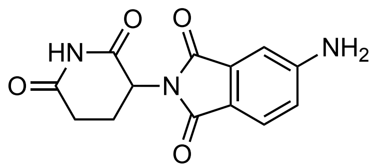

A reported procedure[1] was modified and used to prepare isopomalidomide **4**. 4-aminophthalimide (3.00 g, 18.5 mmol, 1.0 eq) was suspended in $CH_3CN$ (45 mL) and cooled to 0 °C with stirring. A solution of ethyl chloroformate (2.7 mL, 28 mmol, 1.5 eq) in $CH_3CN$ (6 mL) was prepared and added dropwise to the stirring suspension over 60 min and continuously stirred at 0 °C for 1 h. After 1 h, the reaction was warmed to room temperature then stirred for 1 h at room temperature. The resulting brown suspension was concentrated in vacuo to 10 mL. To the suspension was added 0.1% v/v HCl (45 mL) and the suspension was cooled to 0 °C and stirred 30 min. The resulting crystalline product was vacuum filtered over filter paper, washed with a cold mixture of $CH_3CN$/water (1:4, 2 x 5 mL), and dried under high vacuum overnight. The resulting crystalline solid of ethyl 5-amino-1,3-dioxoisoindoline-2-carboxylate was used directly in the following step.

To a dry round bottom flask was added ethyl 5-amino-1,3-dioxoisoindoline-2-carboxylate (2.00 g, 8.54 mmol, 1.0 eq), 3-aminopiperidine-2,6-dione (1.41 g, 8.54 mmol, 1.0 eq), NaOAc (1.40 g, 17.1 mmol, 2.0 eq), and $CH_3CN$ (40 mL) and the suspension was stirred and heated to reflux for 4 h under $N_2$ atmosphere. The reaction was then cooled to room temperature and concentrated in vacuo to 2 mL. To this concentrated mixture was added water (40 mL) and the mixture was stirred at room temperature for 30 min. The resulting crystalline product was vacuum filtered over filter paper, washed with ice cold water (2 x 20 mL) and dried under high vacuum overnight. To a dry round bottom flask was added the dry crystalline product (1.9 g) and DMSO (10 mL). The suspension was heated to 50 °C until completely dissolved. To the clear solution was added acetone (10 mL), then the mixture was allowed to cool to room temperature. To the room temperature mixture was slowly added water (40 mL) over 30 min. The suspension was cooled to 0 °C and stirred for 1 h. The resulting solid was vacuum filtered over filter paper, washed with 5 mL ice cold acetone, and dried under high vacuum overnight to afford **4** (1.55 g, 5.67 mmol, 66%) as a green solid.

**$^1$H NMR** (400 MHz, DMSO-$d_6$) δ 11.05 (s, 1H), 7.52 (d, $J$ = 8.2 Hz, 1H), 6.94 (d, $J$ = 2.0 Hz, 1H), 6.83 (dd, $J$ = 8.3, 2.0 Hz, 1H), 6.55 (s, 2H), 5.01 (dd, $J$ = 12.9, 5.4 Hz, 1H), 2.94 – 2.80 (m, 1H), 2.62 – 2.42 (m, 2H), 2.04 – 1.93 (m, 1H) ppm.

**$^{13}$C NMR** (101 MHz, DMSO-$d_6$) δ 172.8, 170.2, 167.6, 167.1, 155.2, 134.2, 125.2, 116.9, 116.2, 107.0, 48.6, 31.0, 22.2 ppm.

**HRMS** (ESI/LC-TOF) m/z: $[M+H]^+$ Calcd for $C_{13}H_{12}N_3O_4$ 273.0750; Found 273.0743.

**LCMS** (5% → 100% MeCN/$H_2O$, 5 min) $t_R$ = 2.05 min (m/z: $[M+H]^+$ 274).

**3-(4-nitroso-1-oxoisoindolin-2-yl)piperidine-2,6-dione (7)**

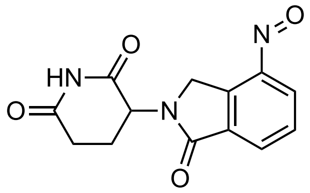


To a solution of lenalidomide (200 mg, 0.77 mmol, 1 eq) in $CH_2Cl_2$ (500 mL) was added a solution of oxone (949 mg, 1.5 mmol, 2.0 eq) in water (3 mL). The biphasic mixture was stirred vigorously at room temperature overnight under $N_2$ atmosphere. The reaction mixture was washed once with water (20 mL), dried over anhydrous $Na_2SO_4$, and concentrated in vacuo by 80%. The resultant green solution of 3-(4-nitroso-1-oxoisoindolin-2-yl)piperidine-2,6-dione (**7**) in $CH_2Cl_2$ was used directly in subsequent steps.

**2-(2,6-dioxopiperidin-3-yl)-5-nitrosoisoindoline-1,3-dione (5)**

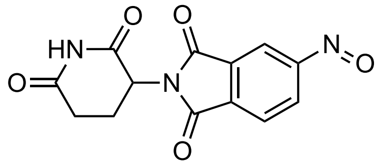

To a round bottom flask was added **4** (101 mg, 0.37 mmol, 1.0 eq) and $CH_2Cl_2$ (40 mL) with stirring. The suspension was gently heated to reflux to promote dissolution, during which additional $CH_2Cl_2$ (210 mL) was added. A portion of the starting material remaining as a suspended solid, the mixture was cooled to room temperature. Oxone (550 mg, 0.90 mmol, 2.4 eq) was dissolved in water (3 mL) and added dropwise to the stirring mixture and stirred vigorously to allow the biphasic reaction to evenly mix. The reaction mixture was stirred at room temperature overnight under $N_2$ atmosphere. The reaction mixture was washed once with water (20 mL), dried over anhydrous $Na_2SO_4$, and concentrated in vacuo by 80%. The resultant green solution of 2-(2,6-dioxopiperidin-3-yl)-5-nitrosoisoindoline-1,3-dione (**5**) in $CH_2Cl_2$ was used directly in subsequent steps.

**3-(4-((4-methoxyphenyl)diazenyl)-1-oxoisoindolin-2-yl)piperidine-2,6-dione (PHOIMiD-1)**

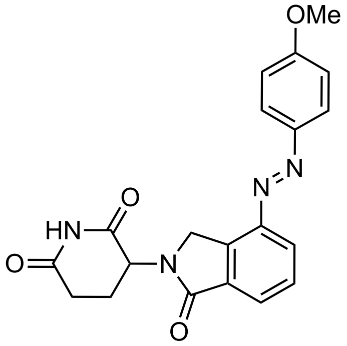


Lenalidomide (50.0 mg, 0.193 mmol, 1.0 eq) was suspended in $CH_2Cl_2$ (50 mL) and ultrasonicated for 3 min. Oxone (237.1 mg, 0.386 mmol, 2.0 eq) was dissolved in water (1.5 mL) and added to the vigorously stirred lenalidomide solution. The reaction was stirred for 18 h, water (5 mL) was added, and the layers were separated. Acetic acid (15 mL) and 4-methoxyaniline (23.8 mg, 0.193 mmol, 1.0 eq) were added to the organic layer and the mixture was concentrated to remove most of the $CH_2Cl_2$. The acetic acid mixture was stirred for 24 h at room temperature and concentrated in vacuo. The crude residue was dissolved with ethyl acetate (25 mL), separated against saturated $NaHCO_3$ (25 mL), washed with 10% LiCl (20 mL) and brine (20 mL), dried over $Na_2SO_4$ and concentrated under reduced pressure. Purification of the resulting crude product by flash column chromatography (0% → 20% MeOH/$CH_2Cl_2$) gave **PHOIMID-1** (49.0 mg, 0.129 mmol, 67%) as a yellow solid.

$\mathbf{R}_f$ = 0.44 (5% MeOH/$CH_2Cl_2$).

**$^1$H NMR** (400 MHz, DMSO-$d_6$) δ 11.02 (s, 1H), 8.22 – 8.15 (m, 1H), 7.99 (d, $J$ = 9.0 Hz, 2H), 7.91 – 7.86 (m, 1H), 7.78 (t, $J$ = 7.7 Hz, 1H), 7.16 (d, $J$ = 9.0 Hz, 2H), 5.22 – 5.12 (m, 1H), 4.96 – 4.63 (m, 2H), 3.89 (s, 3H), 3.00 – 2.86 (m, 1H), 2.69 – 2.55 (m, 2H), 2.09 – 1.98 (m, 1H) ppm.

**$^{13}$C NMR** (101 MHz, DMSO-$d_6$) δ 172.9, 171.0, 167.2, 162.6, 146.6, 146.3, 137.4, 134.3, 133.7, 129.6, 128.4, 124.9, 114.6, 55.8, 51.7, 48.3, 31.2, 22.3 ppm.

**HRMS** (ESI/LC-TOF) m/z: $[M+H]^+$ Calcd for $C_{20}H_{19}N_4O_4$ 379.1401; Found 379.1387.

**LCMS** (5% → 100% MeCN/$H_2O$, 5 min) $t_{R,trans}$ = 3.59 min (m/z: $[M+H]^+$ 379)

$t_{R,cis}$ = 2.84 min (m/z: $[M+H]^+$ 379).

$\boldsymbol{t}_{1/2}$ (37 °C, 1:1 DMSO/PBS) = 17.4 ± 0.15 h.

**3-(4-((4-(morpholinomethyl)phenyl)diazenyl)-1-oxoisoindolin-2-yl)piperidine-2,6-dione (PHOIMiD-2)**

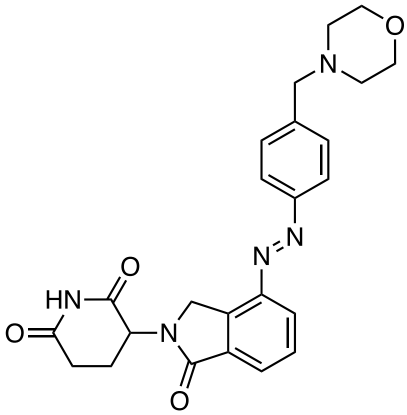

Lenalidomide (30.0 mg, 0.116 mmol, 1.0 eq) was suspended in $CH_2Cl_2$ (30 mL) and ultrasonicated for 3 min. Oxone (142.3 mg, 0.231 mmol, 2.0 eq) was dissolved in water (1 mL) and added to the vigorously stirred lenalidomide solution. The reaction was stirred for 18 h, water (5 mL) was added, and the layers were separated. Acetic acid (10 mL) and 4-morpholin-4-ylmethyl-phenylamine (30.0 mg, 0.116 mmol, 1.0 eq) were added to the organic layer and the mixture was concentrated to remove most of the $CH_2Cl_2$. The acetic acid mixture was stirred for 24 h at room temperature and concentrated in vacuo. The crude residue was dissolved with a 5:1 $CHCl_3$:*i*PrOH mixture (15 mL), separated against saturated $NaHCO_3$ (20 mL), washed with 10% LiCl (15 mL) and brine (15 mL), dried over $Na_2SO_4$ and concentrated under reduced pressure. Purification of the resulting crude product by flash column chromatography (0% → 20% MeOH/$CH_2Cl_2$) gave **PHOIMID-2** (7.9 mg, 0.018 mmol, 15%) as a yellow solid.

$\mathbf{R}_f$ = 0.31 (5% MeOH/$CH_2Cl_2$).

**$^1$H NMR** (400 MHz, DMSO-$d_6$) δ 11.02 (s, 1H), 8.24 (d, *J* = 7.8 Hz, 1H), 8.00 – 7.91 (m, 3H), 7.85 – 7.76 (m, 1H), 7.58 (br s, 2H), 5.18 (dd, *J* = 13.2, 5.1 Hz, 1H), 4.81 (d, *J* = 19.1 Hz, 1H), 4.69 (d, *J* = 19.2 Hz, 1H), 3.61 (br s, 6H), 3.01 – 2.88 (m, 1H), 2.69 – 2.53 (m, 2H), 2.41 (br s, 4H), 2.09 – 1.99 (m, 1H) ppm.

**$^{13}$C NMR** (101 MHz, DMSO-$d_6$) δ 172.9, 171.0, 167.1, 151.2, 146.6, 134.4, 133.8, 131.5, 129.8, 129.7, 129.1, 125.6, 122.7, 115.1, 66.1, 53.1, 51.7, 48.3, 31.2, 22.3 ppm.

**HRMS** (ESI/LC-TOF) m/z: $[M+H]^+$ Calcd for $C_{24}H_{26}N_5O_4$ 448.1979; Found 448.1973.

**LCMS** (5% → 100% MeCN/$H_2O$, 5 min) $t_{R,trans}$ = 2.26 min (m/z: $[M+H]^+$ 448).

$\boldsymbol{t}_{1/2}$ (37 °C, 1:1 DMSO/PBS) = 3.53 ± 0.18 h.

**3-(4-((4-(2-methoxyphenoxy)phenyl)diazenyl)-1-oxoisoindolin-2-yl)piperidine-2,6-dione (PHOIMiD-3)**

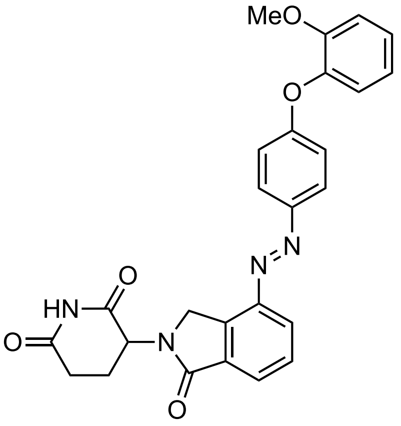


To a solution of **7** (50.0 mg, 0.183 mmol, 1.0 eq) in $CH_2Cl_2$ (25 mL) was added 4-(2-methoxyphenoxy)aniline (39.4 mg, 0.18 mmol, 1.0 eq). The reaction mixture was stirred at room temperature for 10 min. To the stirring reaction mixture was added glacial AcOH (25 mL). The reaction mixture was stirred at room temperature for 62 h, then concentrated in vacuo at 45 °C. The resulting brown residue was dissolved in $CH_2Cl_2$ then washed with saturated aqueous $NaHCO_3$ (100 mL) and saturated aqueous NaCl (100 mL). The organic phase was dried over anhydrous $Na_2SO_4$ and concentrated in vacuo. The crude residue was purified by silica gel flash column chromatography (0% → 1% MeOH/$CH_2Cl_2$). The product fractions were collected and further purified by reverse phase HPLC (5% → 100% $CH_3CN/H_2O$ + 0.1% FA) to afford **PHOIMiD-3** (23.7 mg, 0.050 mmol, 28%) as a yellow solid.

$\mathbf{R}_f$ = 0.49 (10% MeOH/$CH_2Cl_2$).

**$^1$H NMR** (400 MHz, DMSO-*d*$_6$) δ 11.02 (s, 1H), 8.18 (dd, *J* = 7.8, 1.1 Hz, 1H), 7.97 (d, *J* = 9.0 Hz, 2H), 7.89 (d, *J* = 7.2 Hz, 1H), 7.78 (t, *J* = 7.7 Hz, 1H), 7.34 – 7.26 (m, 1H), 7.26 – 7.17 (m, 2H), 7.08 – 7.03 (m, 1H), 7.01 (d, *J* = 9.0 Hz, 2H), 5.17 (dd, *J* = 13.2, 5.1 Hz, 1H), 4.79 (d, *J* = 19.1 Hz, 1H), 4.67 (d, *J* = 19.1 Hz, 1H), 3.75 (s, 3H), 3.00 – 2.87 (m, 1H), 2.66 – 2.46 (m, 2H), 2.09 – 1.99 (m, 1H) ppm.

**$^{13}$C NMR** (101 MHz, DMSO-*d*$_6$) δ 172.9, 171.0, 167.2, 161.2, 151.4, 147.1, 146.6, 142.3, 134.3, 133.7, 129.6, 128.6, 126.6, 125.1, 124.8, 122.3, 121.3, 116.1, 113.6, 55.7, 51.7, 48.2, 31.2, 22.3 ppm.

**HRMS** (ESI/LC-TOF) m/z: $[M+H]^+$ Calcd for $C_{26}H_{23}N_4O_5$ 471.1663; Found 471.1672.

**LCMS** (50% → 100% MeCN/$H_2O$, 5 min) $t_R$ = 4.100 min (m/z: $[M+H]^+$ 471.1).

$\boldsymbol{t}_{1/2}$ (37 °C, DMSO) = 14.3 ± 0.02 h.

$\boldsymbol{t}_{1/2}$ (37 °C, 1:1 DMSO/PBS) = 14.6 ± 0.46 h.

SUPPORTING INFORMATION

**2-(2,6-dioxopiperidin-3-yl)-5-((4-methoxyphenyl)diazenyl)isoindoline-1,3-dione (PHOIMiD-4)**

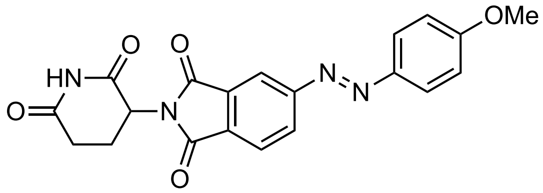


Compound **4** (50.0 mg, 0.183 mmol, 1.0 eq) was suspended in $CH_2Cl_2$ (50 mL) and ultrasonicated for 3 min. Oxone (225 mg, 0.366 mmol, 2.0 eq) was dissolved in water (1.5 mL) and added to the vigorously stirred lenalidomide solution. The reaction was stirred for 18 h, water (5 mL) was added, and the layers were separated. Acetic acid (15 mL) and 4-methoxyaniline (23.8 mg, 0.193 mmol, 1.0 eq) were added to the organic layer and the mixture was concentrated to remove most of the $CH_2Cl_2$. The acetic acid mixture was stirred for 24 h at room temperature and concentrated in vacuo. The crude residue was dissolved with ethyl acetate (25 mL), separated against saturated $NaHCO_3$ (25 mL), washed with 10% LiCl (20 mL) and brine (20 mL), dried over $Na_2SO_4$ and concentrated under reduced pressure. Purification of the resulting crude product by flash column chromatography ($CH_2Cl_2$/MeOH gradient, 0 to 20% MeOH) gave **PHOIMID-4** (9.4 mg, 0.024 mmol, 26%) as a yellow solid.

$\mathbf{R}_f$ = 0.39 (5% MeOH/$CH_2Cl_2$).

**$^1$H NMR** (400 MHz, DMSO-$d_6$) δ 11.16 (s, 1H), 8.31 (dd, *J* = 8.0, 1.7 Hz, 1H), 8.20 (d, *J* = 1.6 Hz, 1H), 8.13 (d, *J* = 7.9 Hz, 1H), 8.02 (d, *J* = 9.0 Hz, 2H), 7.20 (d, *J* = 9.1 Hz, 2H), 5.22 (dd, *J* = 12.9, 5.4 Hz, 1H), 3.91 (s, 3H), 2.97 – 2.85 (m, 1H), 2.68 – 2.53 (m, 2H), 2.16 – 2.06 (m, 1H) ppm.

**$^{13}$C NMR** (101 MHz, DMSO-$d_6$) δ 172.7, 169.8, 166.5, 166.4, 163.2, 156.1, 146.1, 132.8, 131.9, 129.7, 125.5, 125.0, 115.3, 114.9, 55.8, 49.2, 30.9, 22.0 ppm.

**HRMS** (ESI/LC-TOF) m/z: $[M+H]^+$ Calcd for $C_{20}H_{17}N_4O_5$ 393.1193; Found 393.1182.

**LCMS** (50% → 100% MeCN/$H_2O$, 5 min) $t_{R,trans}$ = 3.93 min (m/z: $[M+H]^+$ 393),
$t_{R,cis}$ = 3.26 min (m/z: $[M+H]^+$ 393).

$\boldsymbol{t}_{1/2}$ (37 °C, 1:1 DMSO/PBS) = 14.7 ± 0.12 h.

**2-(2,6-dioxopiperidin-3-yl)-5-((4-isopropylphenyl)diazenyl)isoindoline-1,3-dione (PHOIMiD-5)**

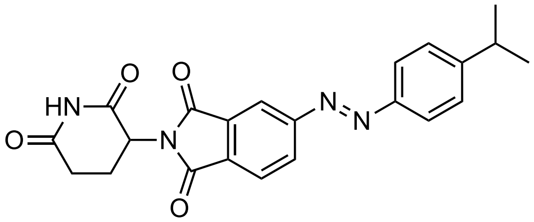

To a solution of **5** (50.0 mg, 0.174 mmol, 1.0 eq) in $CH_2Cl_2$ (30 mL) was added 4-isopropylaniline (28 mg, 0.21 mmol, 1.2 eq). The reaction mixture was stirred at room temperature for 10 min. To the stirring reaction mixture was added glacial AcOH (20 mL). The reaction mixture was stirred at room temperature for 30 h, then concentrated in vacuo at 45 °C. The resulting brown residue was dissolved in $CH_2Cl_2$ then washed with saturated aqueous $NaHCO_3$ (100 mL) and saturated aqueous NaCl (100 mL). The organic phase was dried over anhydrous $Na_2SO_4$ and concentrated in vacuo. The crude residue was purified by reverse phase HPLC (5% → 100% $CH_3CN/H_2O$ + 0.1% FA) to afford **PHOIMiD-5** (19.5 mg, 0.048 mmol, 28%) as a yellow solid.

$\mathbf{R}_f$ = 0.54 (10% MeOH/$CH_2Cl_2$).

**$^1$H NMR** (400 MHz, DMSO-$d_6$) δ 11.16 (s, 1H), 8.33 (dd, $J$ = 7.9, 1.7 Hz, 1H), 8.21 (d, $J$ = 1.1 Hz, 1H), 8.14 (d, $J$ = 7.9 Hz, 1H), 7.94 (d, $J$ = 8.4 Hz, 1H), 7.53 (d, $J$ = 8.5 Hz, 2H), 5.22 (dd, $J$ = 13.0, 5.4 Hz, 1H), 3.03 (hept, $J$ = 6.9 Hz, 1H), 2.96 – 2.84 (m, 1H), 2.68 – 2.53 (m, 2H), 2.15 – 2.06 (m, 1H), 1.27 (d, $J$ = 6.9 Hz, 6H) ppm.

**$^{13}$C NMR** (101 MHz, DMSO-$d_6$) δ 173.2, 170.2, 167.0, 166.8, 156.5, 154.4, 150.6, 133.3, 132.8, 130.4, 128.0, 125.5, 123.9, 115.9, 49.7, 34.0, 31.4, 24.0, 22.4 ppm.

**HRMS** (ESI/LC-TOF) m/z: $[M+H]^+$ Calcd for $C_{22}H_{21}N_4O_4$ 405.1557; Found 405.1562.

**LCMS** (5% → 100% MeCN/$H_2O$, 5 min) $t_R$ = 4.403 min (m/z: $[M+H]^+$ 405.1).

$\boldsymbol{t}_{1/2}$ (37 °C, DMSO) = 1.90 ± 0.002 h.

$\boldsymbol{t}_{1/2}$ (37 °C, 1:1 DMSO/PBS) = 33.6 ± 1.3 h.

**2-(2,6-dioxopiperidin-3-yl)-5-((3-phenoxyphenyl)diazenyl)isoindoline-1,3-dione (PHOIMiD-6)**

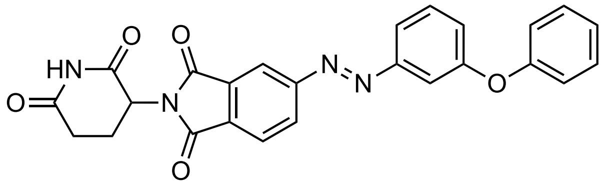


To a solution of **5** (77.5 mg, 0.270 mmol, 1.0 eq) in $CH_2Cl_2$ (47 mL) was added 3-phenoxyaniline (50.0 mg, 0.27 mmol, 1.0 eq). The reaction mixture was stirred at room temperature for 10 min. To the stirring reaction mixture was added glacial AcOH (30 mL). The reaction mixture was stirred at room temperature for 30 h, then concentrated in vacuo at 45 °C. The resulting brown residue was dissolved in $CH_2Cl_2$ then washed with saturated aqueous $NaHCO_3$ (100 mL) and saturated aqueous NaCl (100 mL). The organic phase was dried over anhydrous $Na_2SO_4$ and concentrated in vacuo. The crude residue was purified by silica gel flash column chromatography (0% → 1% MeOH/$CH_2Cl_2$). The product fractions were collected and further purified by reverse phase HPLC (5% → 100% $CH_3CN/H_2O$ + 0.1% FA) to afford **PHOIMiD-6** (20.7 mg, 0.046 mmol, 17%) as a yellow solid.

$\mathbf{R}_f$ = 0.53 (10% MeOH/$CH_2Cl_2$).

$^{1}$**H NMR** (400 MHz, DMSO-$d_6$) δ 11.16 (s, 1H), 8.33 (dd, $J$ = 7.9, 1.7 Hz, 1H), 8.22 (d, $J$ = 1.6 Hz, 1H), 8.13 (d, $J$ = 7.9 Hz, 1H), 7.81 (d, $J$ = 8.4 Hz, 1H), 7.68 (t, $J$ = 8.0 Hz, 1H), 7.51 (t, $J$ = 2.2 Hz, 1H), 7.49 – 7.43 (m, 2H), 7.42 – 7.36 (m, 1H), 7.35 – 7.27 (m, 2H), 7.26 – 7.19 (m, 1H), 7.18 – 7.11 (m, 2H), 7.04 – 6.99 (m, 1H), 5.21 (dd, $J$ = 13.0, 5.4 Hz, 1H), 2.98 – 2.84 (m, 1H), 2.68 – 2.52 (m, 2H), 2.16 – 2.04 (m, 1H) ppm.

$^{13}$**C NMR** (101 MHz, DMSO-$d_6$) δ 172.7, 169.7, 166.4, 166.3, 158.0, 155.9, 155.7, 153.0, 132.8, 132.7, 131.2, 130.3, 130.2, 125.0, 124.2, 122.6, 119.7, 119.3, 115.7, 110.8, 49.3, 30.9, 21.9 ppm.

**HRMS** (ESI/LC-TOF) m/z: $[M+H]^+$ Calcd for $C_{25}H_{19}N_4O_5$ 455.1350; Found 455.1346.

**LCMS** (5% → 100% MeCN/$H_2O$, 5 min) $t_R$ = 4.384 min (m/z: $[M+H]^+$ 455.1).

$\boldsymbol{t}_{1/2}$ (37 °C, DMSO) = 8.54 ± 0.07 h.

$\boldsymbol{t}_{1/2}$ (37 °C, 1:1 DMSO/PBS) = 11.8 ± 0.47 h.

**5-((4-(benzyloxy)phenyl)diazenyl)-2-(2,6-dioxopiperidin-3-yl)isoindoline-1,3-dione (PHOIMiD-7)**

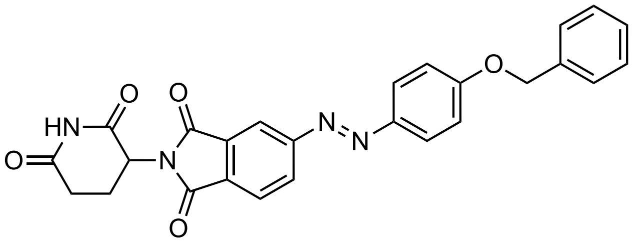

To a solution of **5** (73.1 mg, 0.255 mmol, 1.2 eq) in $CH_2Cl_2$ (37 mL) was added 4-(benzyloxy)aniline hydrochloride (50 mg, 0.21 mmol, 1.0 eq). The reaction mixture was stirred at room temperature for 10 min. To the stirring reaction mixture was added glacial AcOH (30 mL). The reaction mixture was stirred at room temperature for 30 h, then concentrated in vacuo at 45 °C. The resulting brown residue was dissolved in $CH_2Cl_2$ then washed with saturated aqueous $NaHCO_3$ (100 mL) and saturated aqueous NaCl (100 mL). The organic phase was dried over anhydrous $Na_2SO_4$ and concentrated in vacuo. The crude residue was purified by silica gel flash column chromatography (30% → 70% EtOAc/hexanes). The product fractions were collected and further purified by reverse phase HPLC (5% → 100% $CH_3CN/H_2O$ + 0.1% FA) to afford **PHOIMiD-7** (4.1 mg, 0.009 mmol, 4%) as a yellow solid.

$\mathbf{R}_f$ = 0.50 (10% MeOH/$CH_2Cl_2$).

**$^1$H NMR** (400 MHz, DMSO-$d_6$) δ 11.16 (s, 1H), 8.30 (dd, *J* = 7.9, 1.7 Hz, 1H), 8.20 (d, *J* = 1.5 Hz, 1H), 8.12 (d, *J* = 7.8 Hz, 1H), 8.01 (d, *J* = 9.0 Hz, 2H), 7.50 (d, *J* = 7.0 Hz, 2H), 7.45 – 7.40 (m, 2H), 7.39 – 7.34 (m, 1H), 7.28 (d, *J* = 9.0 Hz, 2H), 5.27 (s, 2H), 5.21 (dd, *J* = 13.0, 5.4 Hz, 1H), 2.91 (ddd, *J* = 17.4, 14.0, 5.5 Hz, 1H), 2.69 – 2.53 (m, 2H), 2.14 – 2.06 (m, 1H) ppm.

**$^{13}$C NMR** (101 MHz, DMSO-$d_6$) δ 172.8, 169.8, 166.5, 166.4, 162.2, 156.1, 146.2, 136.4, 132.8, 131.9, 129.7, 128.5, 128.1, 127.9, 125.5, 125.0, 115.7, 115.3, 69.8, 49.2, 30.9, 20.8 ppm.

**HRMS** (ESI/LC-TOF) m/z: $[M+H]^+$ Calcd for $C_{26}H_{21}N_4O_5$ 469.1506; Found 469.1509.

**LCMS** (50% → 100% MeCN/$H_2O$, 5 min) $t_{R,trans}$ = 4.349 min (m/z: $[M+H]^+$ 469.0), $t_{R,cis}$ = 3.763 min (m/z: $[M+H]^+$ 469.1).

$\boldsymbol{t}_{1/2}$ (37 °C, DMSO) = 0.767 ± 0.001 h.

$\boldsymbol{t}_{1/2}$ (37 °C, 1:1 DMSO/PBS) = 7.00 ± 0.09 h.

**5-((4-(2-chlorophenoxy)phenyl)diazenyl)-2-(2,6-dioxopiperidin-3-yl)isoindoline-1,3-dione (PHOIMiD-8)**

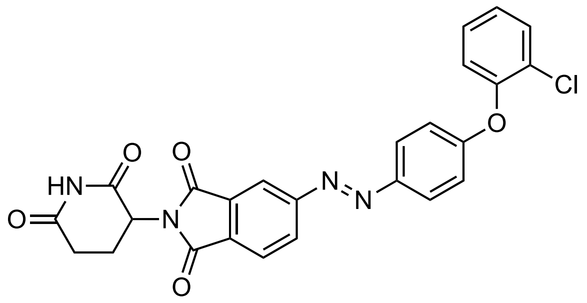

To a solution of **5** (140.0 mg, 0.487 mmol, 1.0 eq) in $CH_2Cl_2$ (80 mL) was added 4-(2-chlorophenoxy)benzenamine (107 mg, 0.49 mmol, 1.0 eq). The reaction mixture was stirred at room temperature for 10 min. To the stirring reaction mixture was added glacial AcOH (50 mL). The reaction mixture was stirred at room temperature for 30 h, then concentrated in vacuo at 45 °C. The resulting brown residue was dissolved in $CH_2Cl_2$ then washed with saturated aqueous $NaHCO_3$ (100 mL) and saturated aqueous NaCl (100 mL). The organic phase was dried over anhydrous $Na_2SO_4$ and concentrated in vacuo. The crude residue was purified by reverse phase HPLC (5% → 100% $CH_3CN/H_2O$ + 0.1% FA) to afford **PHOIMiD-8** (36.1 mg, 0.074 mmol, 15%) as a yellow solid.

$\mathbf{R}_f$ = 0.54 (10% MeOH/$CH_2Cl_2$).

**$^1$H NMR** (400 MHz, DMSO-$d_6$) δ 11.16 (s, 1H), 8.32 (dd, $J$ = 7.9, 1.7 Hz, 1H), 8.21 (d, $J$ = 1.6 Hz, 1H), 8.14 (d, $J$ = 7.9 Hz, 1H), 8.05 (d, $J$ = 9.0 Hz, 2H), 7.72 – 7.64 (m, 1H), 7.53 – 7.43 (m, 1H), 7.40 – 7.31 (m, 2H), 7.14 (d, $J$ = 8.9 Hz, 2H), 5.21 (dd, $J$ = 13.0, 5.4 Hz, 1H), 2.91 (ddd, $J$ = 17.3, 14.0, 5.4 Hz, 1H), 2.68 – 2.51 (m, 2H), 2.16 – 2.04 (m, 1H) ppm.

**$^{13}$C NMR** (101 MHz, DMSO-$d_6$) δ 172.7, 169.7, 166.5, 166.4, 160.5, 156.0, 150.0, 147.5, 132.8, 132.2, 131.0, 129.9, 129.3, 126.9, 125.6, 125.5, 125.0, 122.9, 117.2, 115.4, 49.2, 30.9, 21.9 ppm.

**HRMS** (ESI/LC-TOF) m/z: $[M+H]^+$ Calcd for $C_{25}H_{18}ClN_4O_5$ 491.0943; Found 491.0938.

**LCMS** (5% → 100% MeCN/$H_2O$, 5 min) $t_R$ = 4.518 min (m/z: $[M+H]^+$ 489.0).

$\boldsymbol{t}_{1/2}$ (37 °C, DMSO) = 1.01 ± 0.003 h.

$\boldsymbol{t}_{1/2}$ (37 °C, 1:1 DMSO/PBS) = 8.51 ± 0.16 h.

**2-(2,6-dioxopiperidin-3-yl)-5-((4-(2-methoxyphenoxy)phenyl)diazenyl)isoindoline-1,3-dione (PHOIMiD-9)**

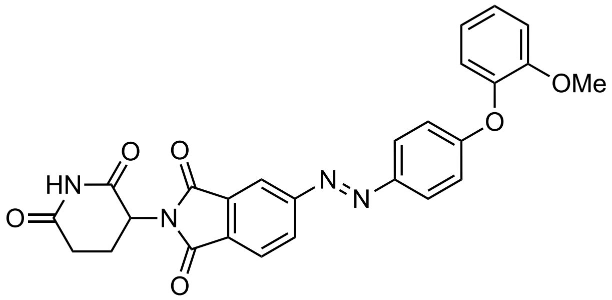


To a solution of **5** (210.3 mg, 0.732 mmol, 1.0 eq) in $CH_2Cl_2$ (125 mL) was added 4-(2-methoxyphenoxy)aniline (157.6 mg, 0.732 mmol, 1.0 eq). The reaction mixture was stirred at room temperature for 10 min. To the stirring reaction mixture was added glacial AcOH (84 mL), resulting in a color change from light yellow to golden after 5 min. The reaction mixture was stirred at room temperature for 90 h, then concentrated in vacuo at 45 °C. The resulting brown residue was dissolved in $CH_2Cl_2$ then washed with saturated aqueous $NaHCO_3$ (100 mL) and saturated aqueous NaCl (100 mL). The organic phase was dried over anhydrous $Na_2SO_4$ and concentrated in vacuo. The crude residue was purified by silica gel flash column chromatography (1% → 3% MeOH/$CH_2Cl_2$). The product fractions were collected and further purified by reverse phase HPLC (5% → 100% $CH_3CN/H_2O$ + 0.1% FA) to afford **PHOIMiD-9** (39.9 mg, 0.082 mmol, 11%) as a yellow solid.

$\mathbf{R}_f$ = 0.51 (10% MeOH/$CH_2Cl_2$).

**$^1$H NMR** (400 MHz, DMSO-$d_6$) δ 11.16 (s, 1H), 8.29 (d, $J$ = 1.7 Hz, 1H), 8.19 (d, $J$ = 1.6 Hz, 1H), 8.13 (d, $J$ = 7.9 Hz, 1H), 8.00 (d, $J$ = 9.0 Hz, 2H), 7.34 – 7.28 (m, 1H), 7.26 – 7.19 (m, 2H), 7.09 – 7.01 (m, 3H), 5.21 (dd, $J$ = 13.0, 5.3 Hz, 1H), 3.75 (s, 3H), 2.91 (ddd, $J$ = 17.3, 14.0, 5.4 Hz, 1H), 2.67 – 2.53 (m, 2H), 2.14 – 2.05 (m, 1H) ppm.

**$^{13}$C NMR** (101 MHz, DMSO-$d_6$) δ 172.8, 169.8, 166.5, 166.4, 161.8, 156.1, 151.4, 146.8, 142.2, 132.8, 132.0, 129.8, 126.8, 125.5, 125.0, 122.5, 121.4, 116.2, 115.3, 113.6, 55.7, 49.2, 30.9, 22.0 ppm.

**HRMS** (ESI/LC-TOF) m/z: $[M + H]^+$ Calcd for $C_{26}H_{21}N_4O_6$ 485.1456; Found 485.1438.

**LCMS** (5% → 100% MeCN/$H_2O$, 5 min) $t_R$ = 1.94 min (m/z: $[M+H]^+$ 485.1).

$\boldsymbol{t}_{1/2}$ (37 °C, DMSO) = 1.02 ± 0.002 h.

$\boldsymbol{t}_{1/2}$ (37 °C, 1:1 DMSO/PBS) = 8.77 ± 0.10 h.

**2-(2,6-dioxopiperidin-3-yl)-5-((4-(3-methoxyphenoxy)phenyl)diazenyl)isoindoline-1,3-dione (PHOIMiD-10)**

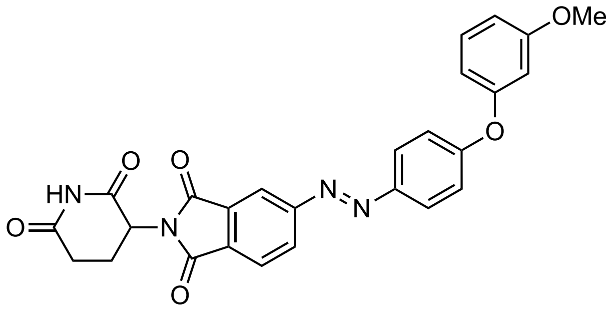


To a solution of **5** (140.0 mg, 0.487 mmol, 1.0 eq) in $CH_2Cl_2$ (80 mL) was added 4-(3-methoxyphenoxy)aniline (104.9 mg, 0.487 mmol, 1.0 eq). The reaction mixture was stirred at room temperature for 10 min. To the stirring reaction mixture was added glacial AcOH (50 mL). The reaction mixture was stirred at room temperature for 30 h, then concentrated in vacuo at 45 °C. The resulting brown residue was dissolved in $CH_2Cl_2$ then washed with saturated aqueous $NaHCO_3$ (100 mL) and saturated aqueous NaCl (100 mL). The organic phase was dried over anhydrous $Na_2SO_4$ and concentrated in vacuo. The crude residue was purified by reverse phase HPLC (5% → 100% $CH_3CN/H_2O$ + 0.1% FA) to afford **PHOIMiD-10** (43.0 mg, 0.089, 18%) as a yellow solid.

$\mathbf{R}_f$ = 0.53 (10% MeOH/$CH_2Cl_2$).

**$^1$H NMR** (400 MHz, DMSO-$d_6$) δ 11.17 (s, 1H), 8.33 (dd, $J$ = 7.9, 1.7 Hz, 1H), 8.22 (d, $J$ = 1.4 Hz, 1H), 8.14 (d, $J$ = 8.0 Hz, 1H), 8.06 (d, $J$ = 8.9 Hz, 2H), 7.39 (t, $J$ = 8.2 Hz, 1H), 7.22 (d, $J$ = 9.0 Hz, 2H), 6.88 – 6.82 (m, 1H), 6.79 – 6.71 (m, 2H), 5.22 (dd, $J$ = 13.0, 5.3 Hz, 1H), 3.78 (s, 3H), 2.92 (ddd, $J$ = 17.4, 14.0, 5.4 Hz, 1H), 2.68 – 2.54 (m, 2H), 2.15 – 2.07 (m, 1H) ppm.

**$^{13}$C NMR** (101 MHz, DMSO-$d_6$) δ 173.2, 170.2, 167.0, 166.8, 161.4, 161.4, 156.6, 156.5, 147.9, 133.3, 132.7, 131.3, 130.3, 126.1, 125.5, 118.7, 115.9, 112.3, 111.1, 106.4, 55.9, 49.7, 31.4, 22.4 ppm.

**HRMS** (ESI/LC-TOF) m/z: $[M+Na]^+$ Calcd for $C_{26}H_{20}N_4NaO_6$ 507.1275; Found 507.1264.

**LCMS** (5% → 100% MeCN/$H_2O$, 5 min) $t_R$ = 4.430 min (m/z: $[M+H]^+$ 485.1).

$\boldsymbol{t}_{1/2}$ (37 °C, DMSO) = 50.5 ± 0.12 h.

$\boldsymbol{t}_{1/2}$ (37 °C, 1:1 DMSO/PBS) = 12.4 ± 0.25 h.

SUPPORTING INFORMATION

**2-(2,6-dioxopiperidin-3-yl)-5-((4-(4-methoxyphenoxy)phenyl)diazenyl)isoindoline-1,3-dione (PHOIMiD-11)**

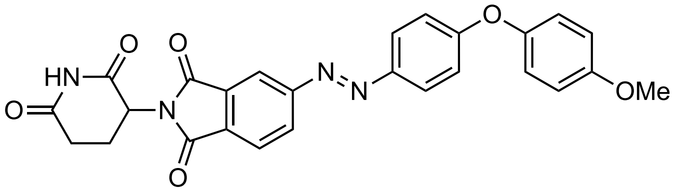

To a solution of **5** (140.0 mg, 0.487 mmol, 1.0 eq) in $CH_2Cl_2$ (80 mL) was added 4-(4-methoxyphenoxy)benzenamine (105 mg, 0.49 mmol, 1.0 eq). The reaction mixture was stirred at room temperature for 10 min. To the stirring reaction mixture was added glacial AcOH (50 mL). The reaction mixture was stirred at room temperature for 30 h, then concentrated in vacuo at 45 °C. The resulting brown residue was dissolved in $CH_2Cl_2$ then washed with saturated aqueous $NaHCO_3$ (100 mL) and saturated aqueous NaCl (100 mL). The organic phase was dried over anhydrous $Na_2SO_4$ and concentrated in vacuo. The crude residue was purified by reverse phase HPLC (5% → 100% $CH_3CN/H_2O$ + 0.1% FA) to afford **PHOIMiD-11** (43.7 mg, 0.090, 19%) as a yellow solid.

$\mathbf{R}_f$ = 0.53 (10% MeOH/$CH_2Cl_2$).

**$^1$H NMR** (400 MHz, DMSO-$d_6$) δ 11.16 (s, 1H), 8.31 (dd, $J$ = 7.9, 1.7 Hz, 1H), 8.20 (d, $J$ = 1.6 Hz, 1H), 8.13 (d, $J$ = 7.9 Hz, 1H), 8.02 (d, $J$ = 9.0 Hz, 2H), 7.19 – 7.08 (m, 3H), 7.04 (d, $J$ = 9.1 Hz, 2H), 5.21 (dd, $J$ = 13.0, 5.3 Hz, 1H), 3.79 (s, 3H), 2.91 (ddd, $J$ = 17.4, 14.0, 5.4 Hz, 1H), 2.67 – 2.53 (m, 2H), 2.16 – 2.05 (m, 1H) ppm.

**$^{13}$C NMR** (101 MHz, DMSO-$d_6$) δ 172.7, 169.7, 166.5, 166.4, 162.2, 156.5, 156.0, 147.9, 147.0, 132.8, 132.1, 129.8, 125.6, 125.0, 121.7, 117.2, 115.3, 115.3, 55.5, 49.2, 30.9, 22.0 ppm.

**HRMS** (ESI/LC-TOF) m/z: $[M+Na]^+$ Calcd for $C_{26}H_{20}N_4NaO_6$ 507.1275; Found 507.1278.

**LCMS** (5% → 100% MeCN/$H_2O$, 5 min) $t_R$ = 4.330 min (m/z: $[M+H]^+$ 485.1).

$\boldsymbol{t}_{1/2}$ (37 °C, DMSO) = 40.4 ± 0.13 h.

$\boldsymbol{t}_{1/2}$ (37 °C, 1:1 DMSO/PBS) = 6.29 ± 0.10 h.

##### III. UV-Vis Spectroscopy

UV-Vis spectra were recorded on a Cary 60 Scan UV-Vis spectrophotometer (Agilent) equipped with an 18-cell holder. Samples were prepared under red light in a dark room by diluting compound stock solutions (10 mM in DMSO) to a final concentration of 25 μM with DMSO or DMSO/PBS. Samples were measured in UV-transparent plastic micro cuvettes with a 10 mm pathlength (BrandTech).

For wavelengths scans and photocycling experiments, photoswitching was carried out using an OptoScan monochromator (Cairn) powered by an OptoSource Illuminator (Cairn) equipped with a 75 mW xenon arc lamp (Ushio). The monochromator was connected using BNC connections to a data acquisition (DAQ) device (National Instruments), which was controlled using a PC with control programs written in MATLAB (MathWorks). Samples were irradiated with light using a fiber optic cable connected to the monochromator. For wavelength scans, a UV-Vis absorbance spectrum (200-800 nm) was recorded in the dark, followed by irradiation from 370 nm to 600 nm in 5 minute steps with a spectrum recorded at the end of each 5 minute period (supplementary Table S1). For photocycling experiments, absorbance was measured at 370 nm, starting with a 10 minute dark period. Next, the sample was irradiated, alternating between 390 nm and 500 nm light with 5 minutes of irradiation per cycle. A total of 36 full cycles (10 minutes each) were recorded.

For thermal relaxation experiments, samples were sonicated for 10 minutes and tapped carefully on the benchtop to displace bubbles. Samples were loaded into the cell holder and equilibrated to 37 °C. Absorbance at 370 nm or 380 nm was measured for each sample in quick succession using the Kinetics program (Cary WinUV) at a rate of one cycle per minute. Experiments started with a 10 minute dark period to establish and record a fully relaxed baseline. Next, samples were irradiated with 390 nm light using an LED plate for 1 h to fully switch compounds. Lastly, full dark conditions were restored and samples were carefully sealed to prevent evaporation during the measurement of thermal relaxation for 10–24 h. Data were exported as CSV files and plotted using Prism 9 (GraphPad). Thermal relaxation half-life values were determined by fitting the data to an exponential one-phase decay model in Prism 9.

**Table S1**. UV-Vis wavelength scan steps.

| Step | Wavelength (nm) | Time (s) |
|---|---|---|
| 1[a] | 590 | 20 |
| 2 | 370 | 300 |
| 3 | 380 | 300 |
| 4 | 390 | 300 |
| 5 | 400 | 300 |
| 6 | 410 | 300 |
| 7 | 420 | 300 |
| 8 | 430 | 300 |
| 9 | 440 | 300 |
| 10 | 450 | 300 |
| 11 | 460 | 300 |
| 12 | 470 | 300 |
| 13 | 480 | 300 |
| 14 | 490 | 300 |
| 15 | 500 | 300 |
| 16 | 510 | 300 |
| 17 | 520 | 300 |
| 18 | 530 | 300 |
| 19 | 540 | 300 |
| 20 | 550 | 300 |
| 21 | 560 | 300 |
| 22 | 570 | 300 |
| 23 | 580 | 300 |
| 24 | 590 | 300 |
| 25 | 600 | 300 |

[a] The light path was blocked during the first step allowing a dark spectrum to be recorded.

#### IV. Cell Lines and Reagents

Human acute lymphoblastic leukemia (RS4;11) cells (ATCC CRL-1873) were cultured in full growth medium consisting of phenol red-free RPMI1640 medium (Gibco) supplemented with 10% FBS (Gibco), 100 U/mL penicillin (Gibco), and 100 µg/mL streptomycin (Gibco), and were maintained in a humidified incubator at 37 °C with 5% $CO_2$ in air. U2OS cells stably expressing the ZF constructs were cultured in Dulbecco’s modified Eagle’s medium (DMEM) (ThermoFisher Scientific) supplemented with 10% FBS (ThermoFisher Scientific), 100 U/mL Antibiotic-Antimycotic (ThermoFisher Scientific), and 1 µg/mL puromycin (ThermoFisher Scientific). The 14 lentiviral ZF plasmids were generated using the Cilantro 2 degradation reporter vector (Addgene, 74450) as previously described.[2] The amino acid sequences of the 14 ZFs are listed in supplementary Table S2.

#### V. Automated Imaging Screening and Analysis

PhenoPlate™ 96-well microplates (PerkinElmer, 055302) were printed with 10 mM of stock compounds in varying volumes using Tecan D300e Digital Dispenser (Tecan) and HP T8+ dispense head cassettes (F0L59A). U2OS cells stably expressing the ZF degron reporters were seeded in 100 µL of 10000 cells per well in compound preprinted 96 well plates. The assay plates containing the cells and test compounds were incubated in light-proof boxes in the dark or with 390 nm pulsed irradiation (100 ms / 10 s; 1% duty cycle). After 24 hours of incubation, the 96-well plates were washed once with PBS then fixed in 4% paraformaldehyde and stained by Hoechst 33342 (ThermoFisher Scientific, H3570). Imaging of eGFP and mCherry was performed for each plate using an Opera Phenix imaging system followed by analysis with Harmony Software v4.9 (PerkinElmer). The fluorescence intensity of eGFP was normalized to that of mCherry for every cell. The mean normalized eGFP intensity of all cells in each well was then normalized by that in DMSO-treated wells to determine the GFP level relative to DMSO for each compound across doses. The GFP level relative to DMSO was used to generate a heatmap using GraphPad Prism 9.

**Table S2**. Amino acid sequences of the zinc finger motifs cloned into the cilantro 2 vector.

| Protein (residue numbers) | Amino acid sequence |
|---|---|
| ZNF276 (524-546 aa) | LQCEVCGFQCRQRASLKYHMTKH |
| ZNF653 (556-578 aa) | LQCEICGYQCRQRASLNWHMKKH |
| ZNF827 (374-396 aa) | FQCPICGLVIKRKSYWKRHMVIH |
| ZFP91 (400-422 aa) | LQCEICGFTCRQKASLNWHMKKH |
| E4F1 (220-242 aa) | HECKLCGASFRTKGSLIRHHRRH |
| ZNF787 (178-200 aa) | FVCPRCGRGFSQPKSLARHLRLH |
| ZNF517 (452-474 aa) | YRCRACGRACSRLSTLIQHQKVH |
| ZNF654 (25-47 aa) | FACVICGRKFRNRGLMQKHLKNH |
| PATZ1 (383-405 aa) | YSCPVCGLRFKRKDRMSYHVRSH |
| ZNF582 (395-417 aa) | YQCKVCGRAFKRVSHLTVHYRIH |
| ZKSC5 (430-452 aa) | YGCNECGKNFGRHSHLIEHLKRH |
| IKZF3 (146-168 aa) | FQCNQCGASFTQKGNLLRHIKLH |
| IKZF3 Q147E (146-168 aa) | FECNQCGASFTQKGNLLRHIKLH |
| SALL4 (410-432 aa) | FVCSVCGHRFTTKGNLKVHFHRH |

#### VI. Cell Proliferation Assays

All steps were performed under strict dark conditions with red light for visibility during preparation. Compound stock solutions (10 mM in DMSO) were serially diluted to 2X final concentrations with phenol red-free full growth medium. RS4;11 cells in phenol red-free full growth medium (100 µL) were seeded at 15,000 cells per well into 96-well plates. Cells were treated with 2X PHOIMiD stocks (100 µL) in triplicate for a final concentration of 100 µM to 300 nM with 1% DMSO. In addition, each assay plate was prepared with blanks containing full growth medium (1% DMSO) and negative controls containing cells treated with full growth medium (1% DMSO) and no compound. The assay plates were incubated in light-proof boxes in the dark or with pulsed 390 nm irradiation (100 ms / 10 s; 1% duty cycle). After 72 h, 10 µL of CellTiter 96® $AQ_{ueous}$ One Solution Reagent (Promega) was added to each well and incubated for 5-7 h in the dark at 37 °C. The absorbance at 500 nm was measured using a FLUOstar Omega plate reader (BMG Labtech). Data were blank corrected by subtracting the average blank absorbance and normalized to the average negative control absorbance. Dose-response curve fitting and $IC_{50}$ quantification were performed using the '[Inhibitor] vs. response -- Variable slope (four parameters)' fit in Prism 9 (GraphPad). Data in figures are the mean viability ± SEM.

#### VII. Immunoblotting

All steps were performed under strict dark conditions with red light for visibility during preparation. Compound stock solutions (10 mM in DMSO) were serially diluted to 2X final concentrations with phenol red-free full growth medium. RS4;11 cells in phenol red-free full growth medium (2 mL) were seeded at $2x10^6$ cells per well into 6-well plates. The cells were treated with 2X stocks (2 mL) to reach the final treatment concentrations with 1% DMSO in each condition. The cells were incubated in light-proof boxes in the dark or with pulsed 390 nm irradiation (100 ms / 10 s; 1% duty cycle) for the indicated times. After incubation, cells were collected in the dark by centrifugation (200 x *g*, 5 min, 4 °C) and washed twice with ice-cold PBS (1 mL). Cell pellets were immediately lysed or stored at –80 °C. Cells were lysed in RIPA lysis buffer (Thermo Scientific) supplemented with protease inhibitor cocktail (Roche) and phosphatase inhibitor cocktail (Sigma-Aldrich) on ice for 20 minutes. The lysates were centrifuged at 15,000 x g for 10 minutes at 4 °C. The supernatant was transferred to a new microcentrifuge tube and kept on ice. The protein concentrations were determined using the BCA assay (Pierce). The samples were supplemented with Laemmli buffer (Bio-Rad) containing 2-mercaptoethanol (355 mM; Sigma-Aldrich) and denatured at 95 °C for 10 minutes and immediately resolved by SDS-PAGE or stored at –20 °C. Equal amounts of lysate were loaded onto a 4–12% Bis-Tris gel (Invitrogen) and resolved by SDS-PAGE. Proteins were transferred to a PVDF membrane (Millipore), stained with Ponceau S Staining Solution (Cell Signaling Technology) or Pierce Reversible Protein Stain Kit (Thermo Scientific), and blocked with 5% Blotting Grade Blocker Non Fat Dry Milk (Bio-Rad) in TBS with 0.1% Tween-20 (TBST) for 30 minutes at room temperature. The membranes were incubated with primary antibody overnight at 4 °C and washed four times with TBST (Table S2). Next, the membranes were incubated with HRPconjugated secondary antibodies for 30 minutes at room temperature then washed four times with TBST (Table S2). The membranes were incubated for 5 minutes with SuperSignal West Pico PLUS or SuperSignal West Femto Maximum Sensitivity Substrate (Thermo Scientific) and the chemiluminescent signal was acquired using a ChemiDoc imaging system (Bio-Rad) or an ImageQuant LAS 400 (GE).

**Table S3**. Antibody information and concentrations.

| Antibody | Dilution Factor (WB) | Source | Identifier |
|---|---|---|---|
| anti-Rabbit IgG, peroxidase-linked species-specific whole antibody (from donkey) Secondary Antibody | 1:5000 | Thermo Fisher Scientific | NA934V |
| anti-Mouse IgG, peroxidase-linked species-specific whole antibody (from sheep) Secondary Antibody | 1:5000 | Thermo Fisher Scientific | NA931V |
| Rabbit anti-Vinculin | 1:2000 | Bethyl Laboratories | A302-535A |
| Ikaros (D6N9Y) Rabbit mAb | 1:1000 | Cell Signaling Technology | #14859 |
| Aiolos (D1C1E) Rabbit mAb | 1:1000 | Cell Signaling Technology | #15103 |
| Proliferating Cell Nuclear Antigen, PC10, Unconjugated, Culture supernatant | 1:2000 | Agilent Dako | M0879 |

#### VIII. Supplemental Figures

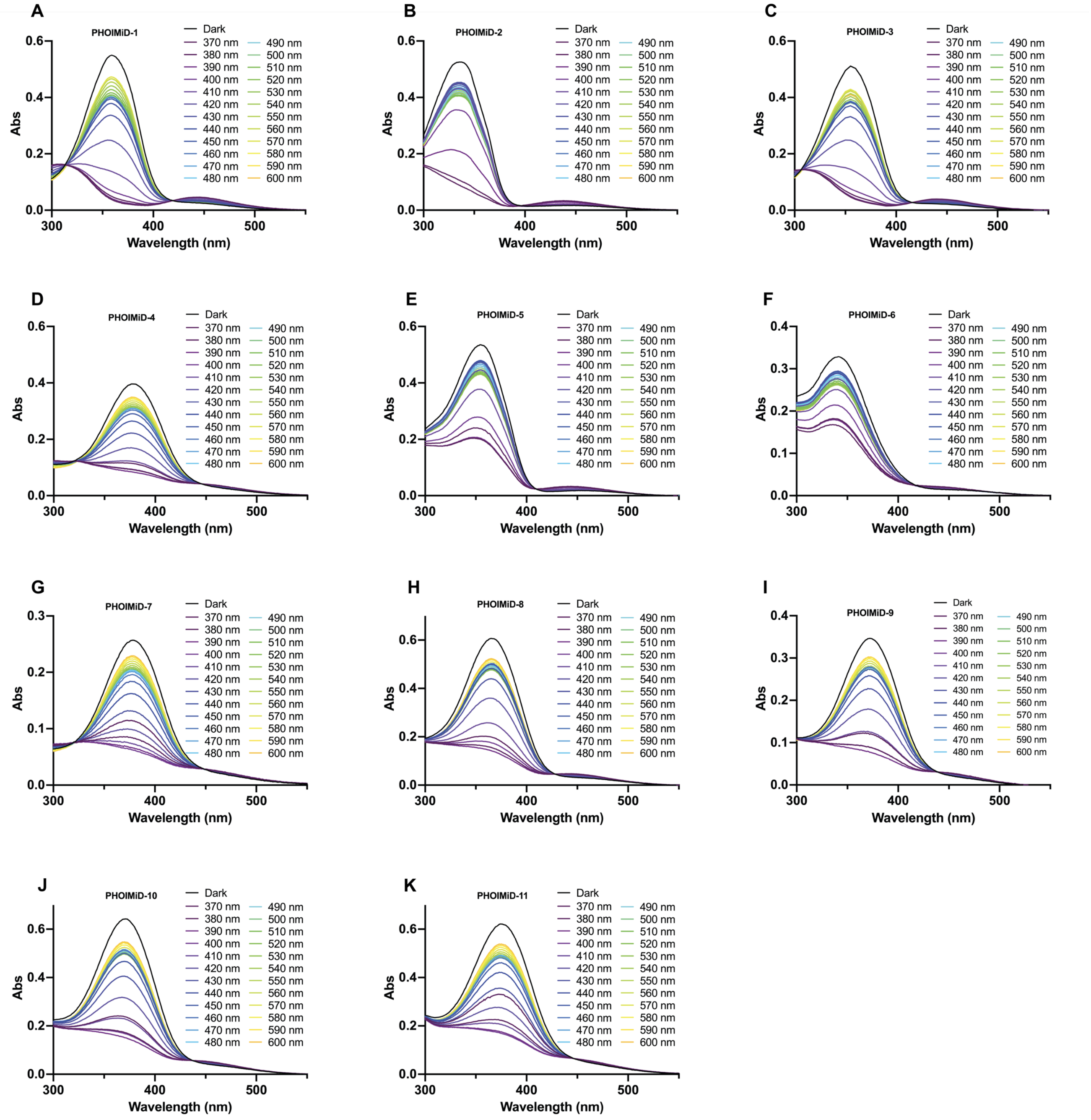


**Figure S1**. UV-Vis wavelength scans for **PHOIMiD-1–11**.

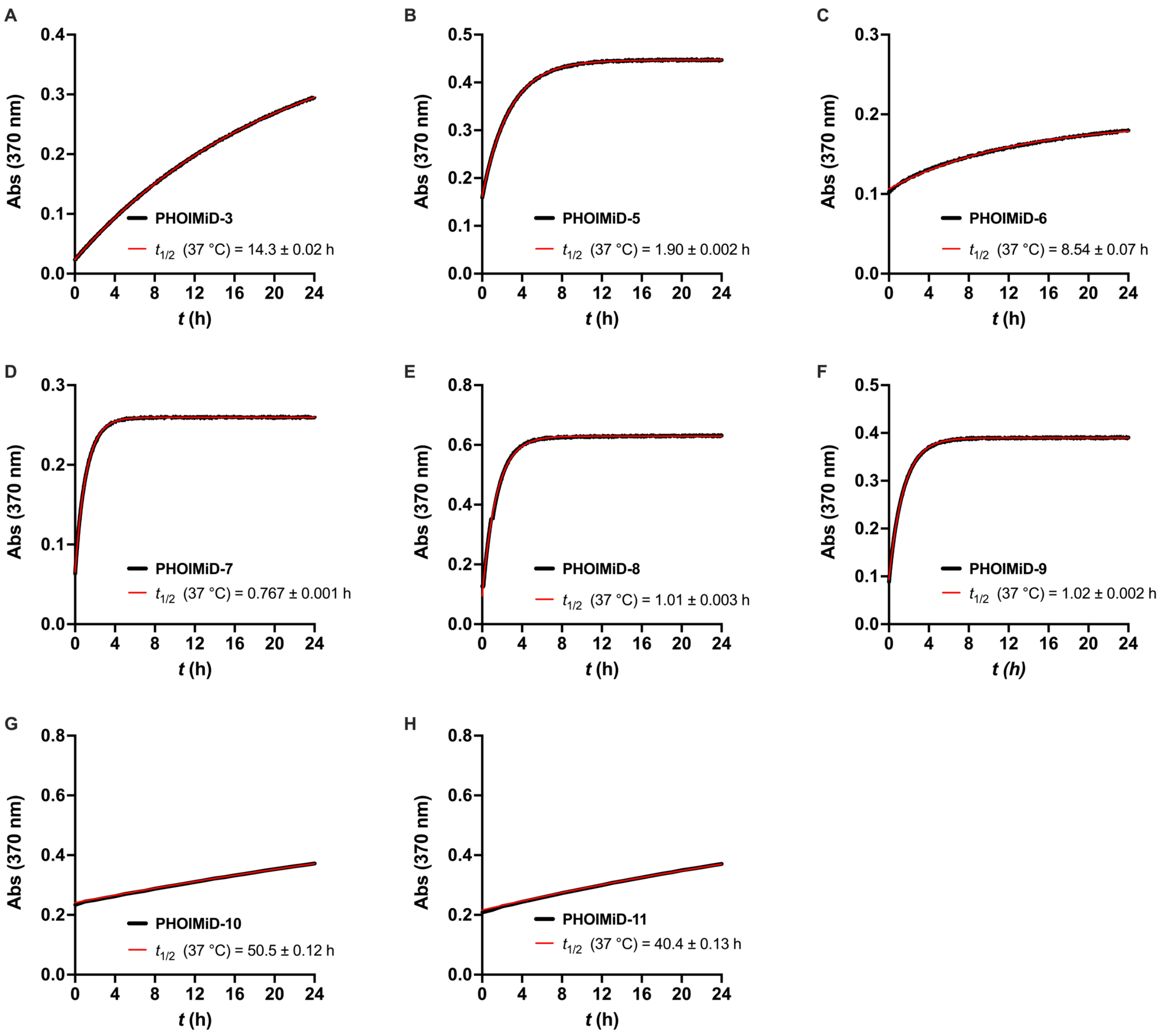


**Figure S2**. Thermal relaxation of PHOIMiDs in DMSO at 37 °C.

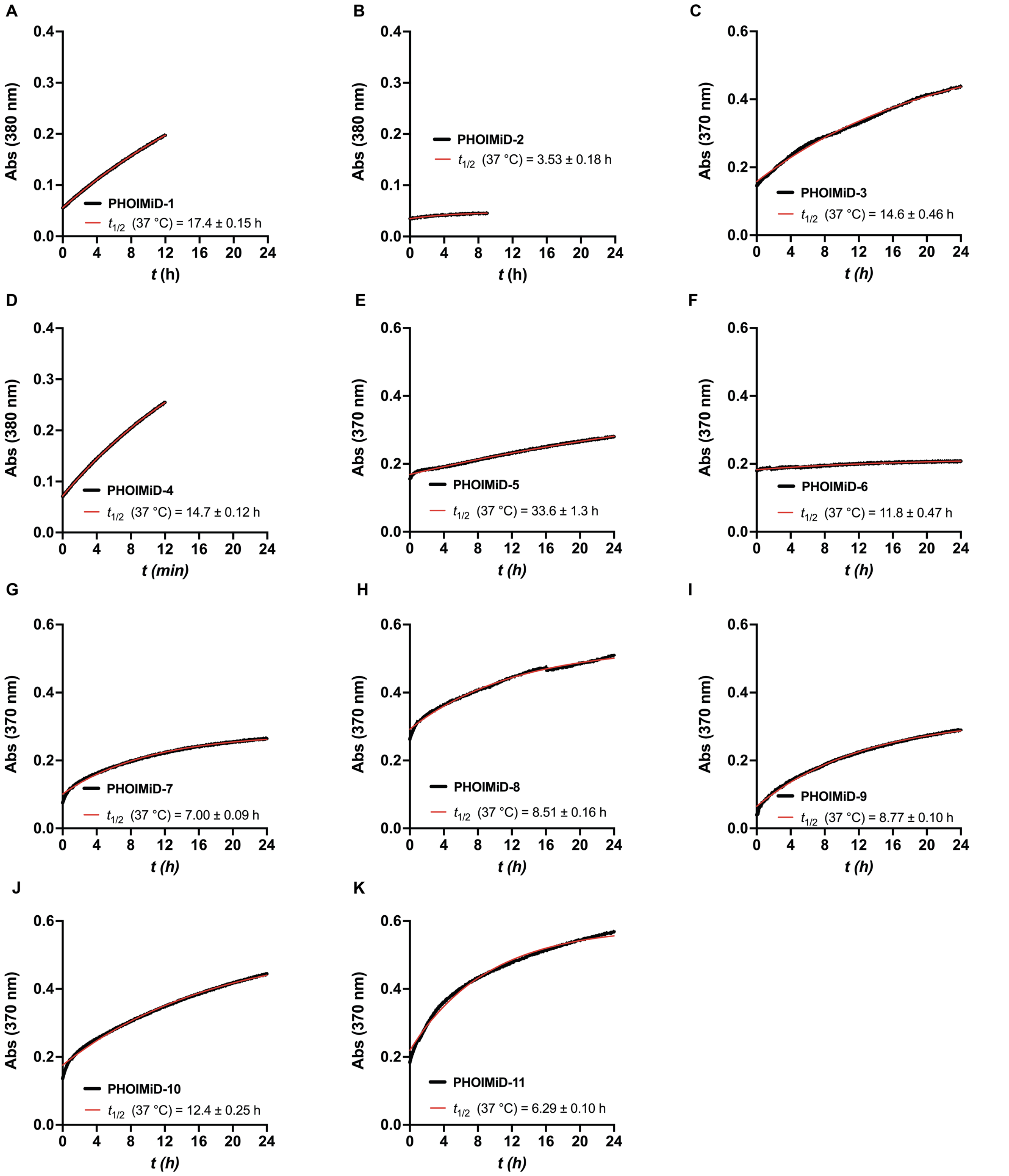


**Figure S3**. Thermal relaxation of PHOIMiDs in 50% DMSO/PBS at 37 °C.

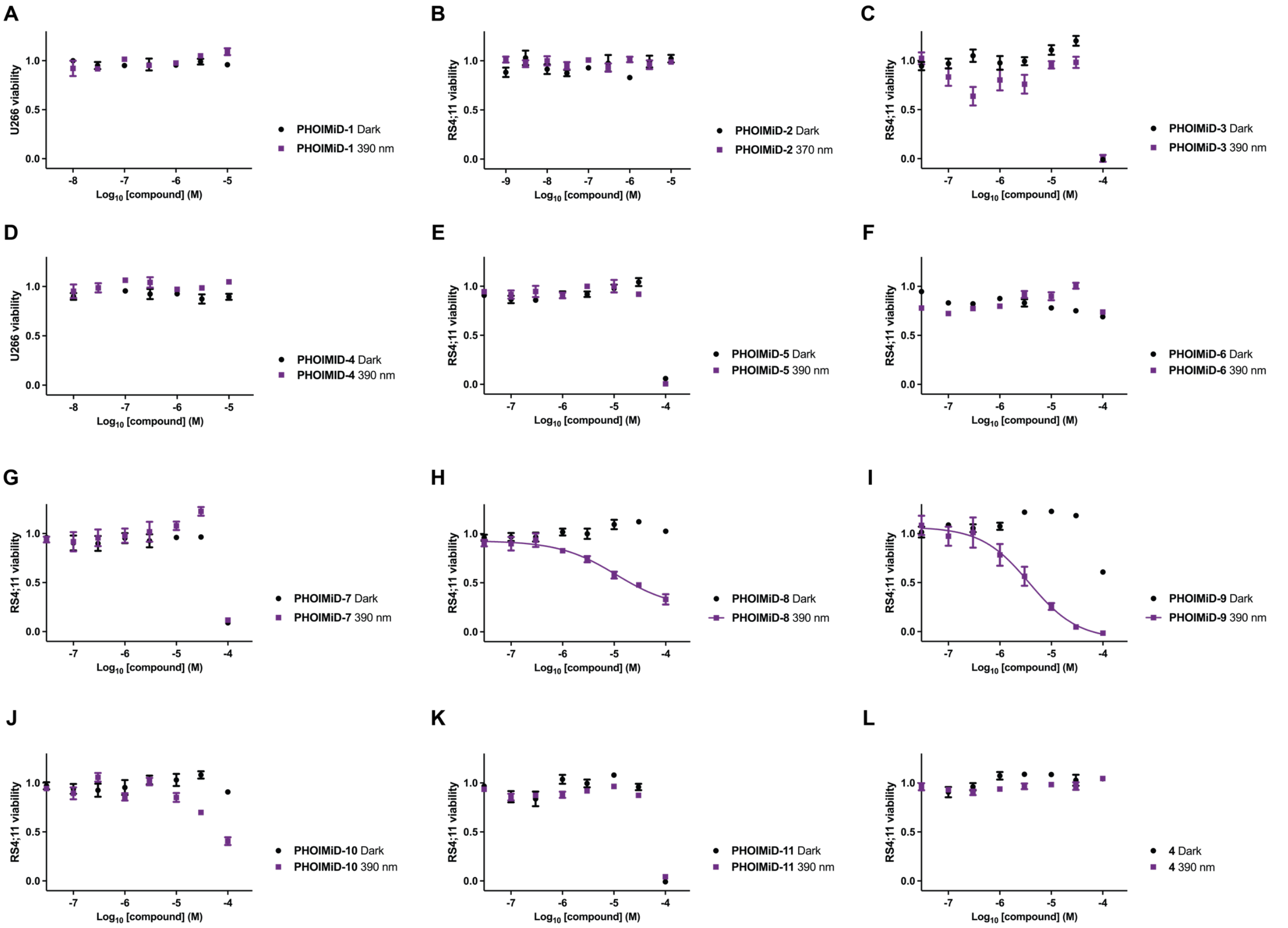


**Figure S4**. MTS data for **PHOIMiD-1–11** and isopomalidomide (**4**)**.** The cells were treated at the indicated concentrations under pulsed 390 nm irradiation (100 ms per 10 s) or in the dark.

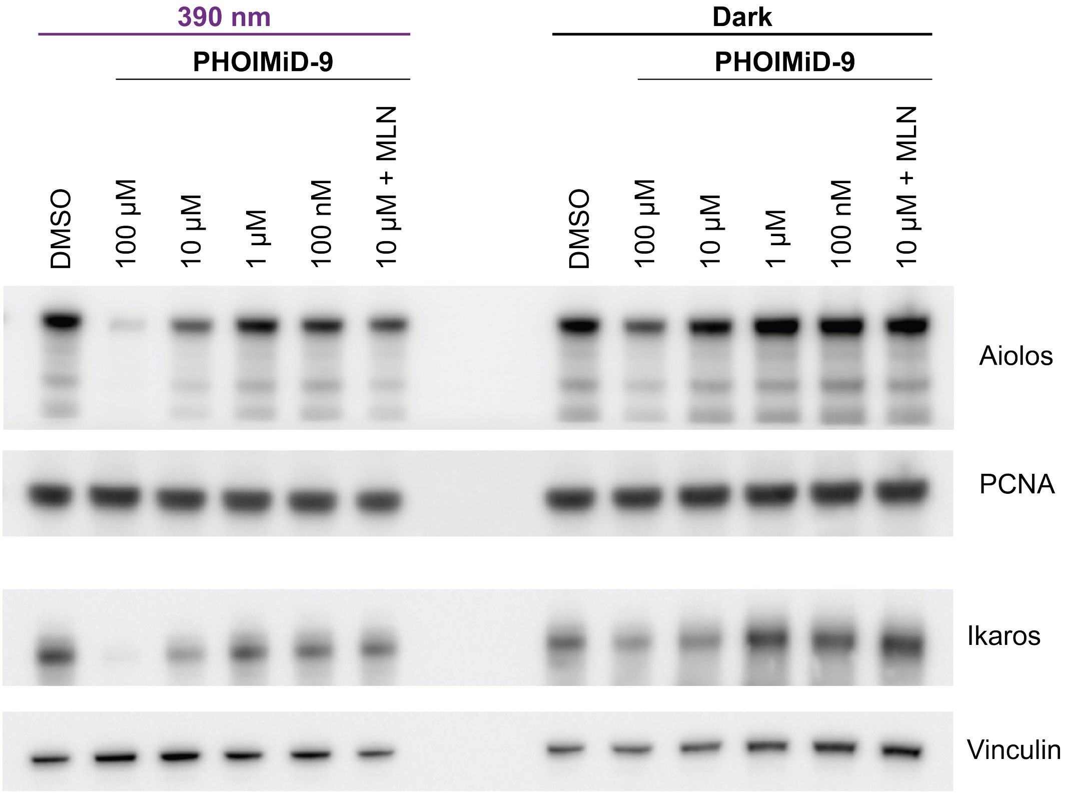


**Figure S5**. Western blot of Ikaros and Aiolos in U266 cells treated for 24 h with **PHOIMiD-9** at the indicated concentrations under pulsed 390 nm irradiation (100 ms per 10 s) or in the dark. MLN (1 µM MLN4924) was used as an additional control.

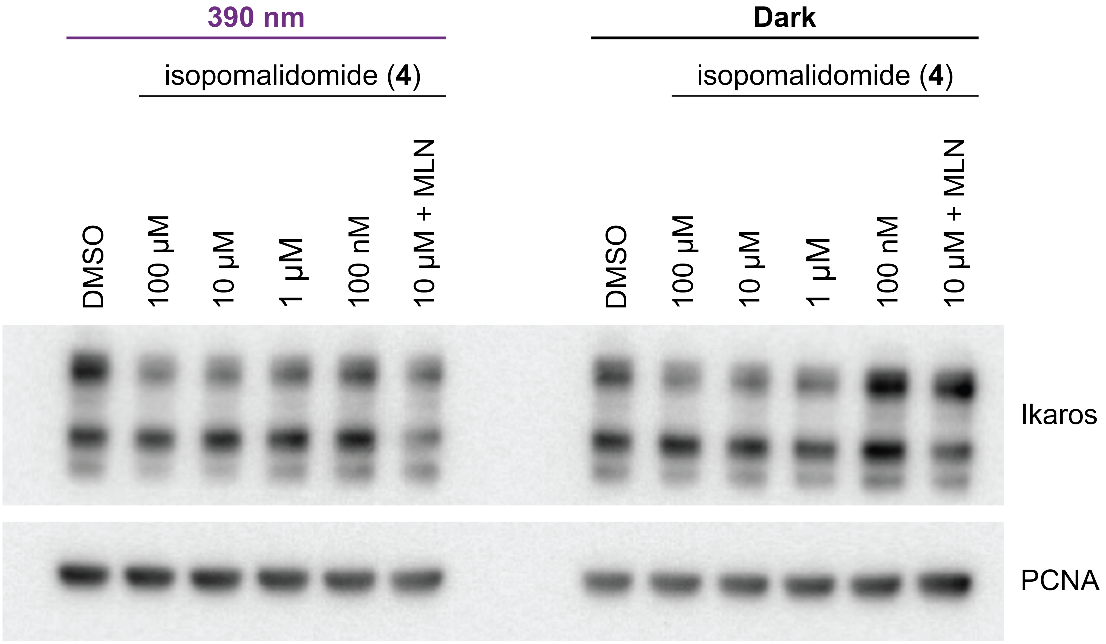


**Figure S6**. Western blot of Ikaros in RS4;11 cells treated for 24 h with **4** at the indicated concentrations under pulsed 390 nm irradiation (100 ms per 10 s) or in the dark. MLN (1 µM MLN4924) was used as an additional control.

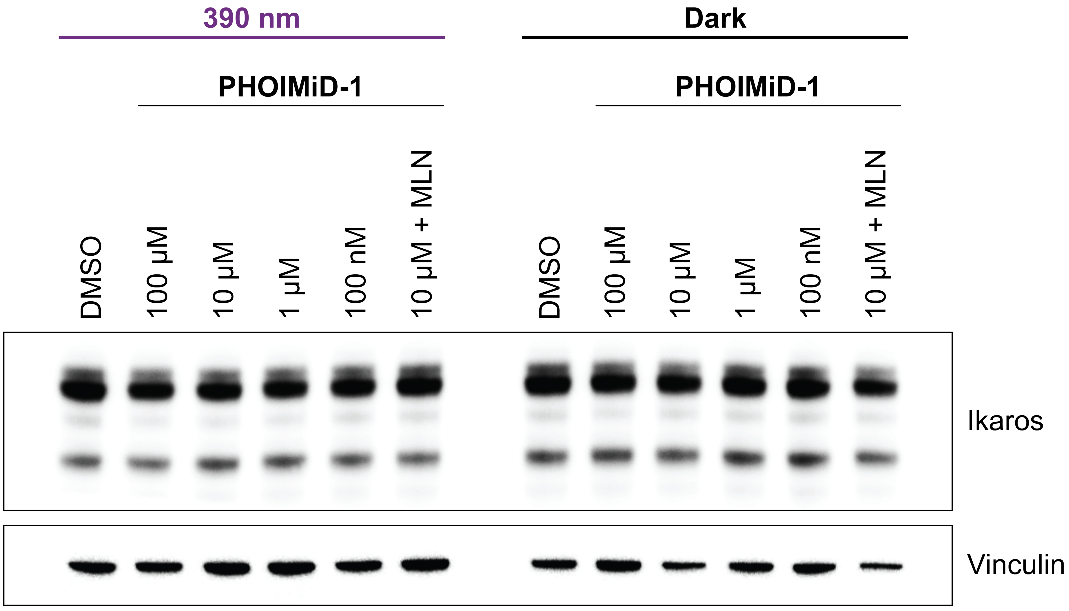


**Figure S7**. Western blot of Ikaros in RS4;11 cells treated for 12 h with **PHOIMiD-1** at the indicated concentrations under pulsed 390 nm irradiation (100 ms per 10 s) or in the dark. MLN (1 µM MLN4924) was used as an additional control.

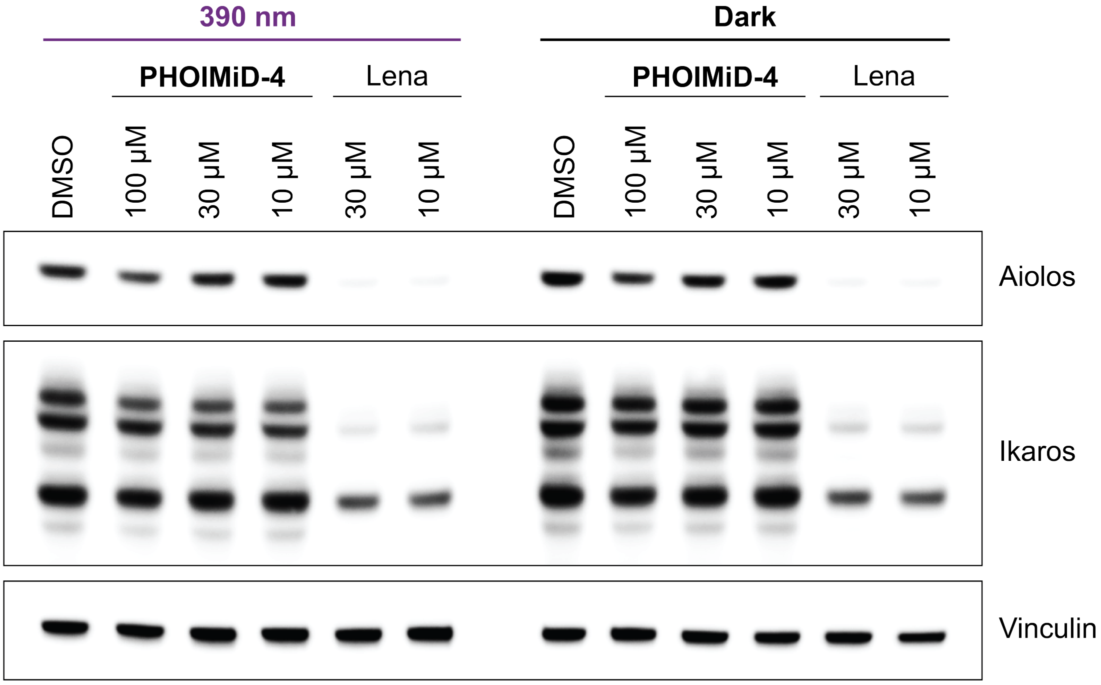


**Figure S8**. Western blot of RS4;11 cells treated with **PHOIMiD-4** or lenalidomide (Lena) for 12 h at the indicated concentrations under pulsed 390 nm irradiation (100 ms per 10 s) or in the dark.

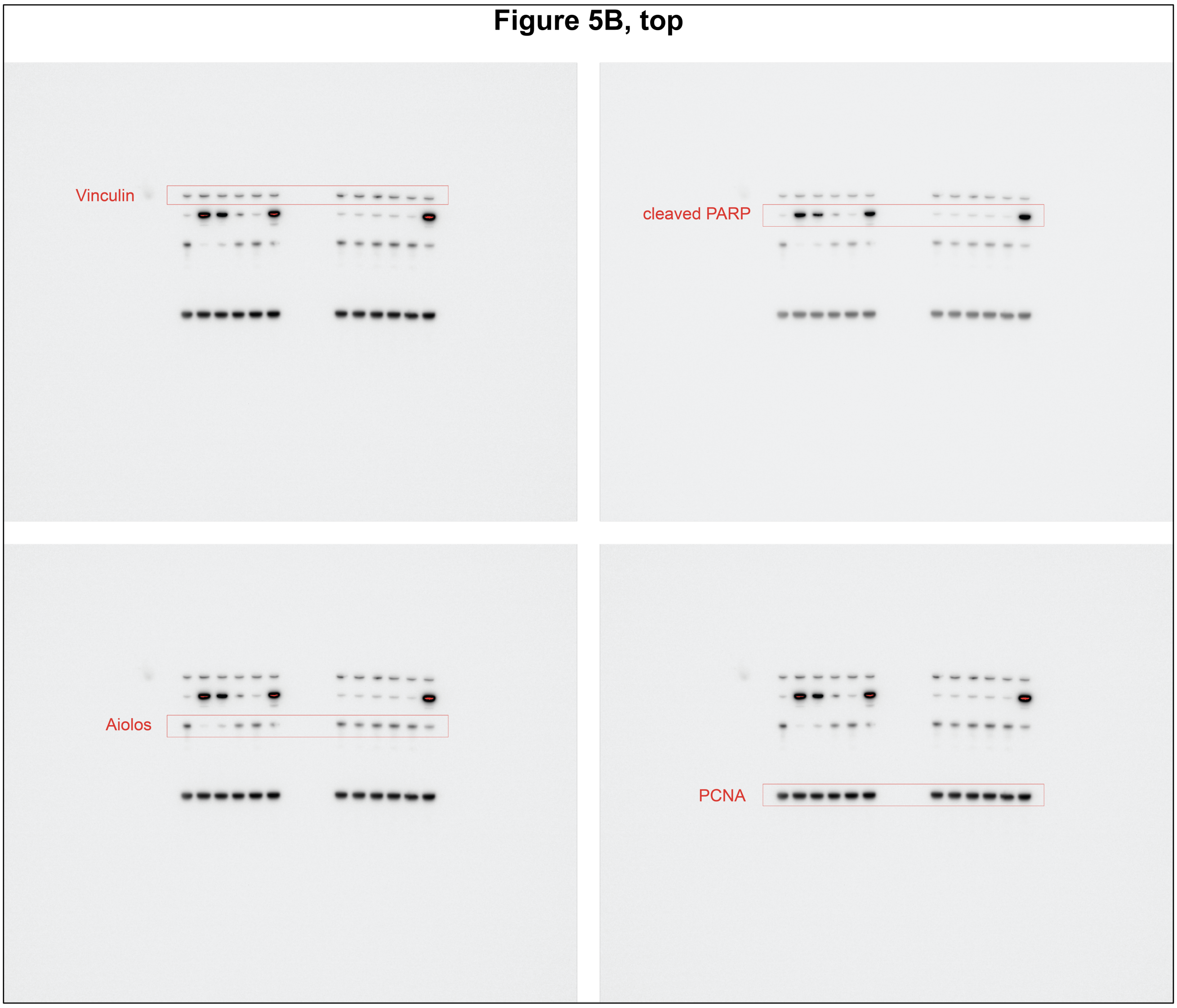


**Figure S9**. Uncropped Western blots for Figure 5B, top.

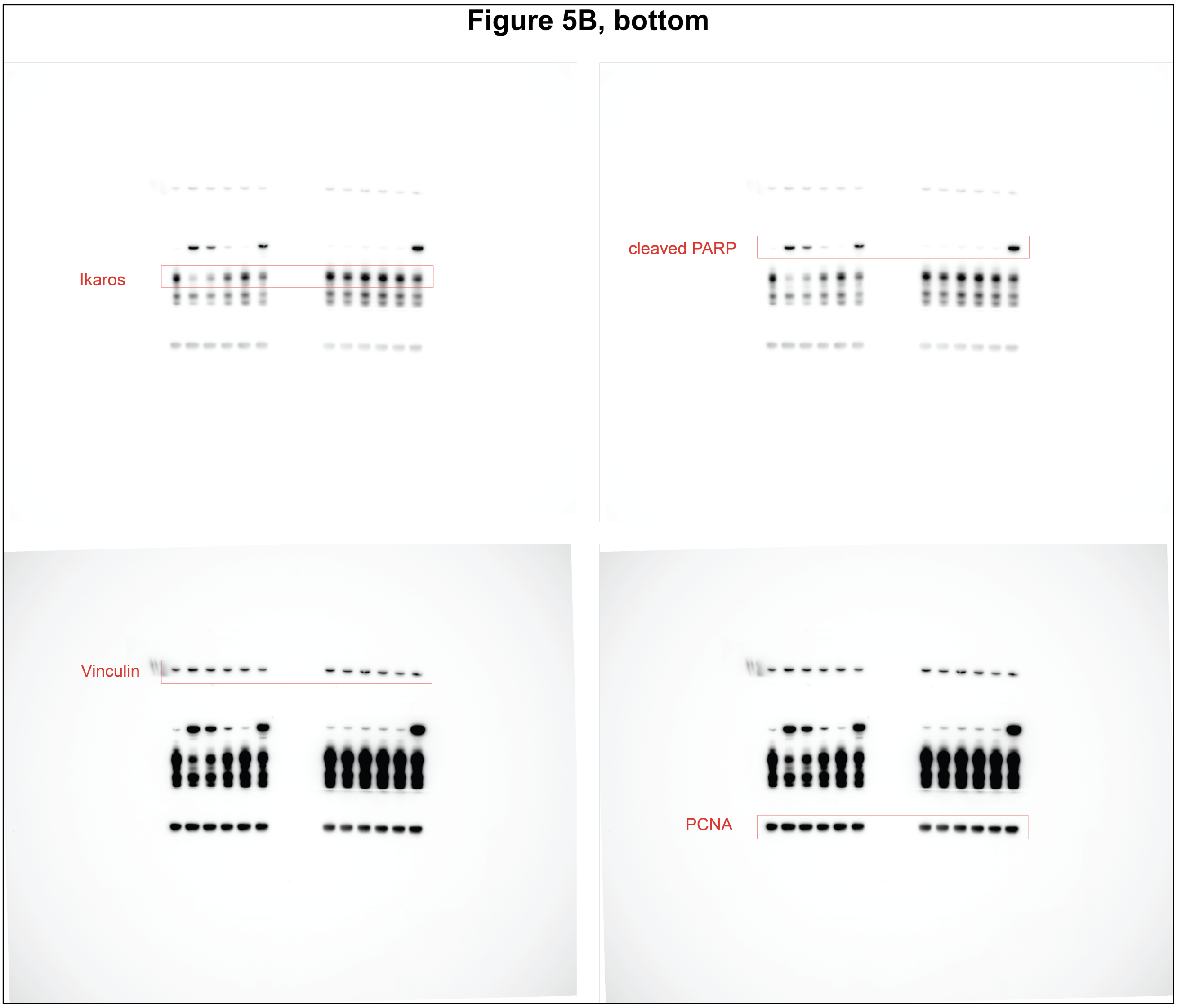


**Figure S10**. Uncropped Western blots for Figure 5B, bottom.

**Figure 5C**

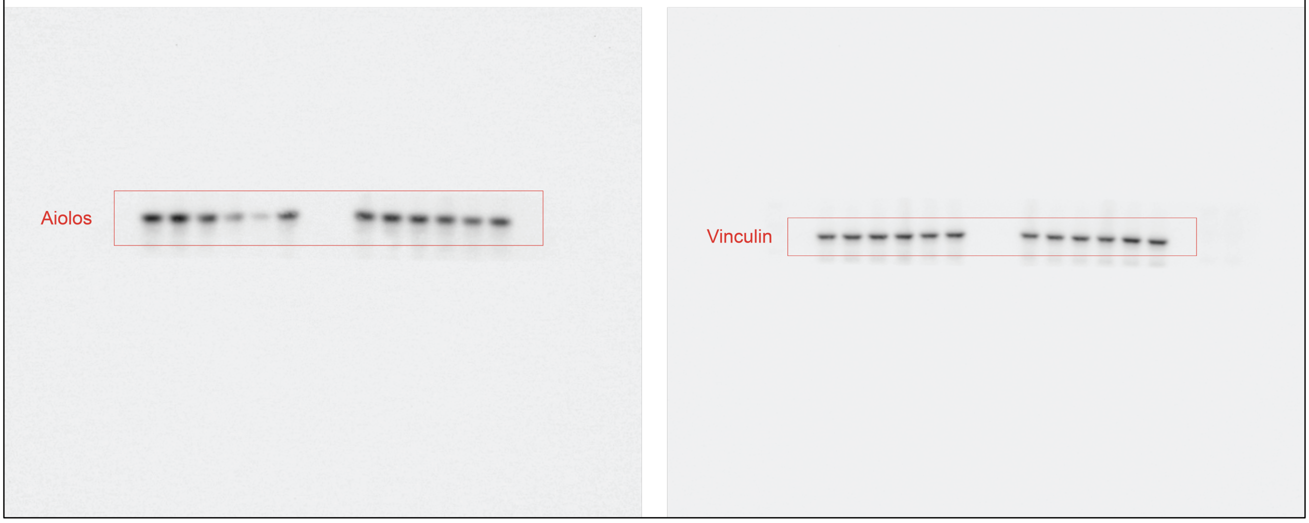


**Figure S11**. Uncropped Western blots for Figure 5C.

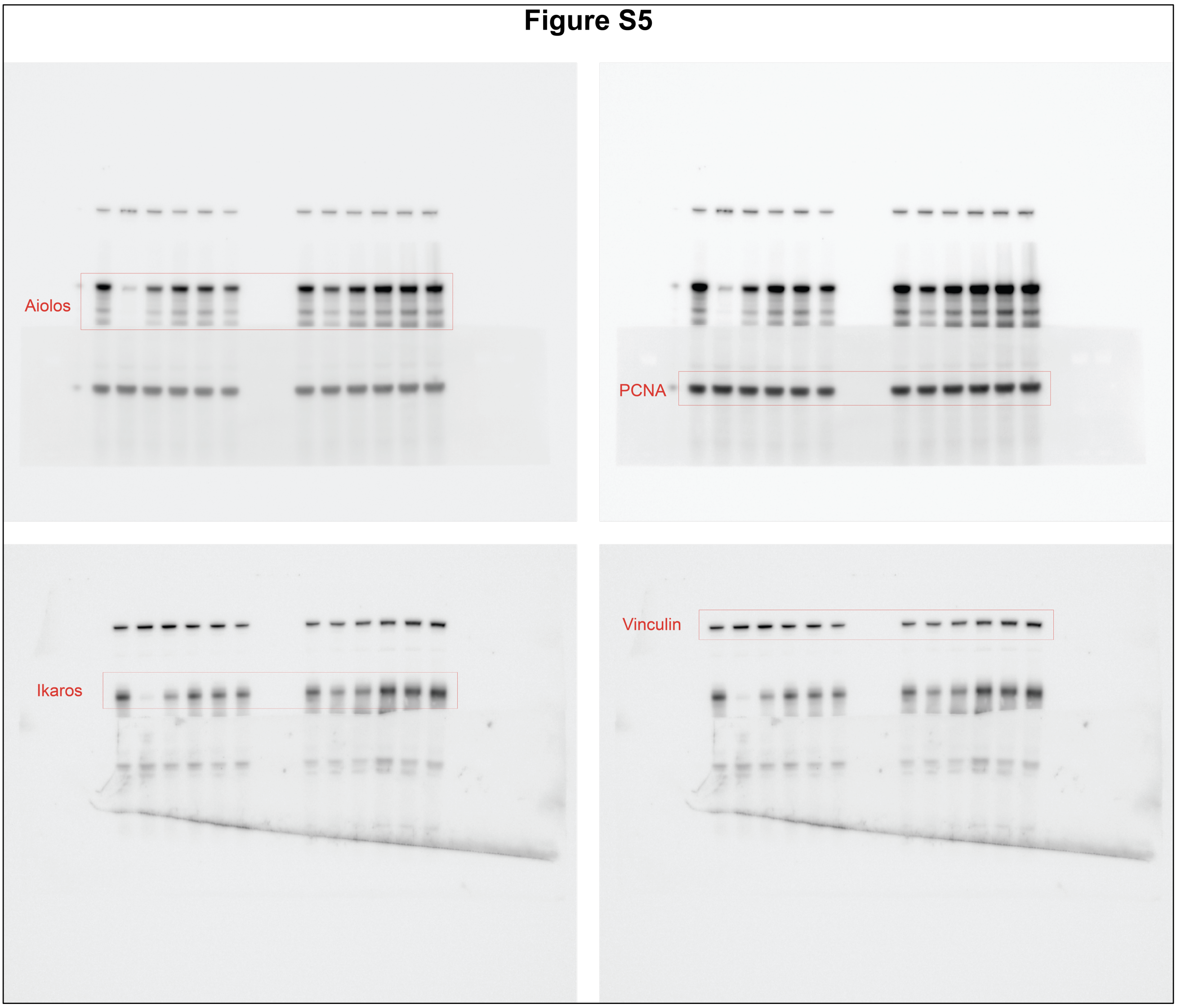


**Figure S12**. Uncropped Western blots for Figure S5.

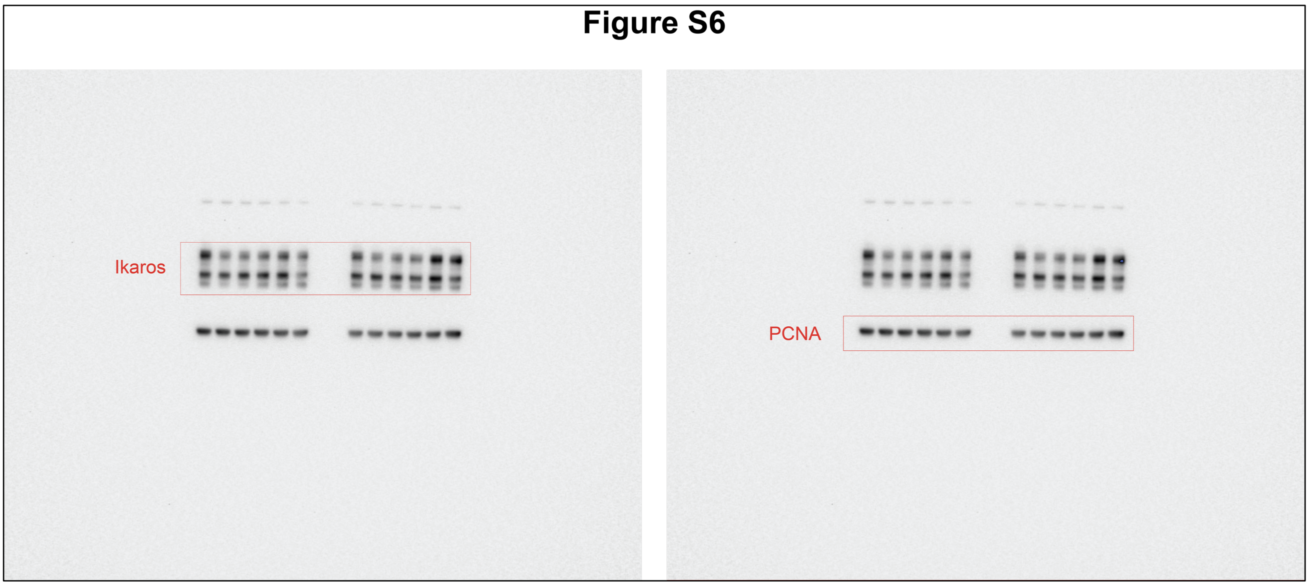


**Figure S13**. Uncropped Western blots for Figure S6.

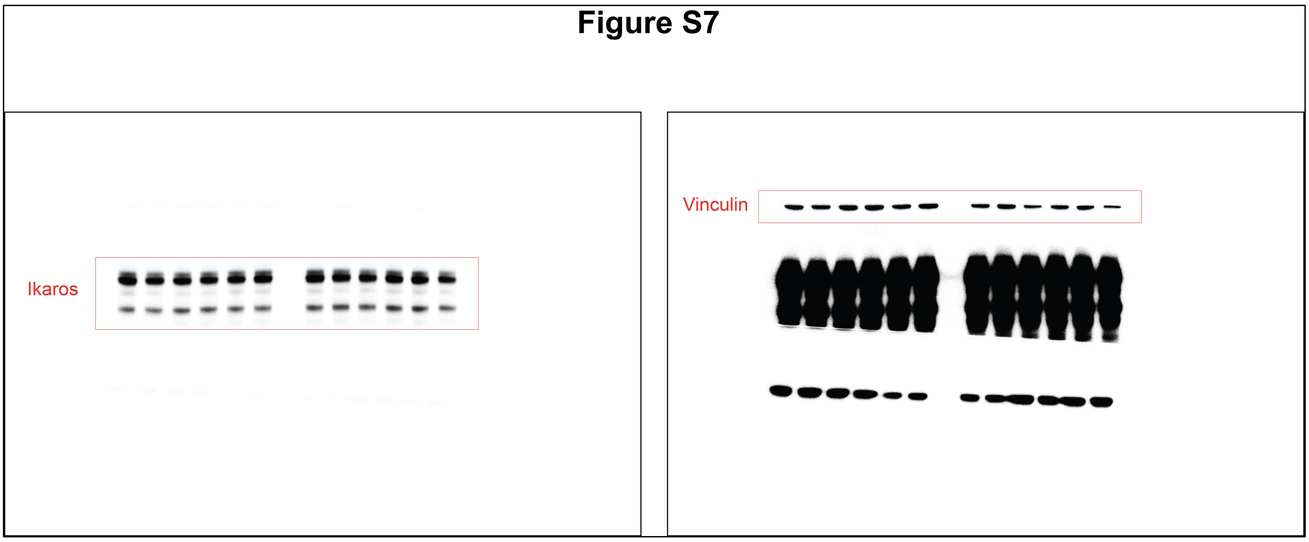


**Figure S14**. Uncropped Western blots for Figure S7.

#### Figure S8

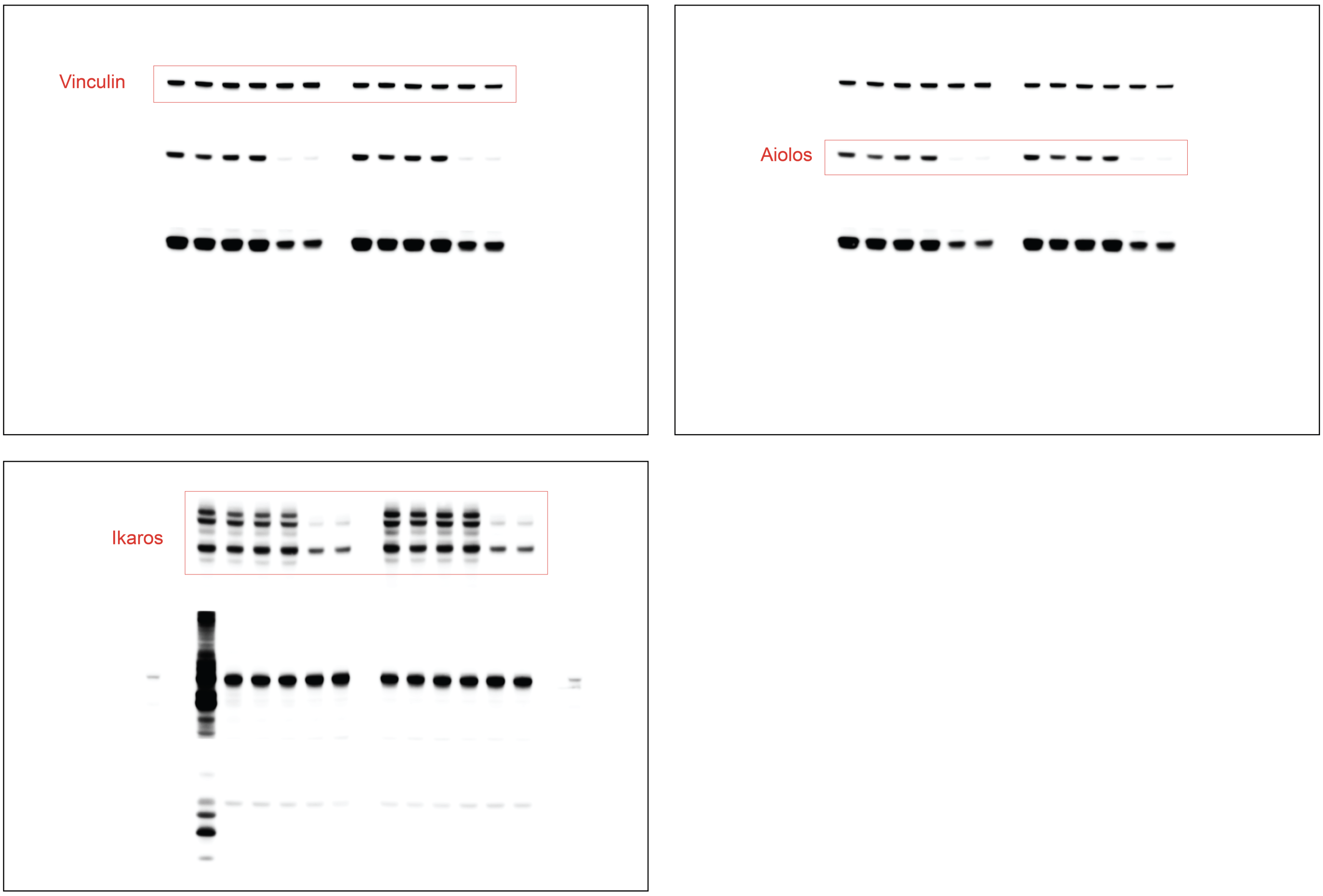


**Figure S15**. Uncropped Western blots for Figure S8.

#### X. NMR Spectra

### SUPPORTING INFORMATION

$^{1}$H NMR (400 MHz, DMSO-$d_6$)

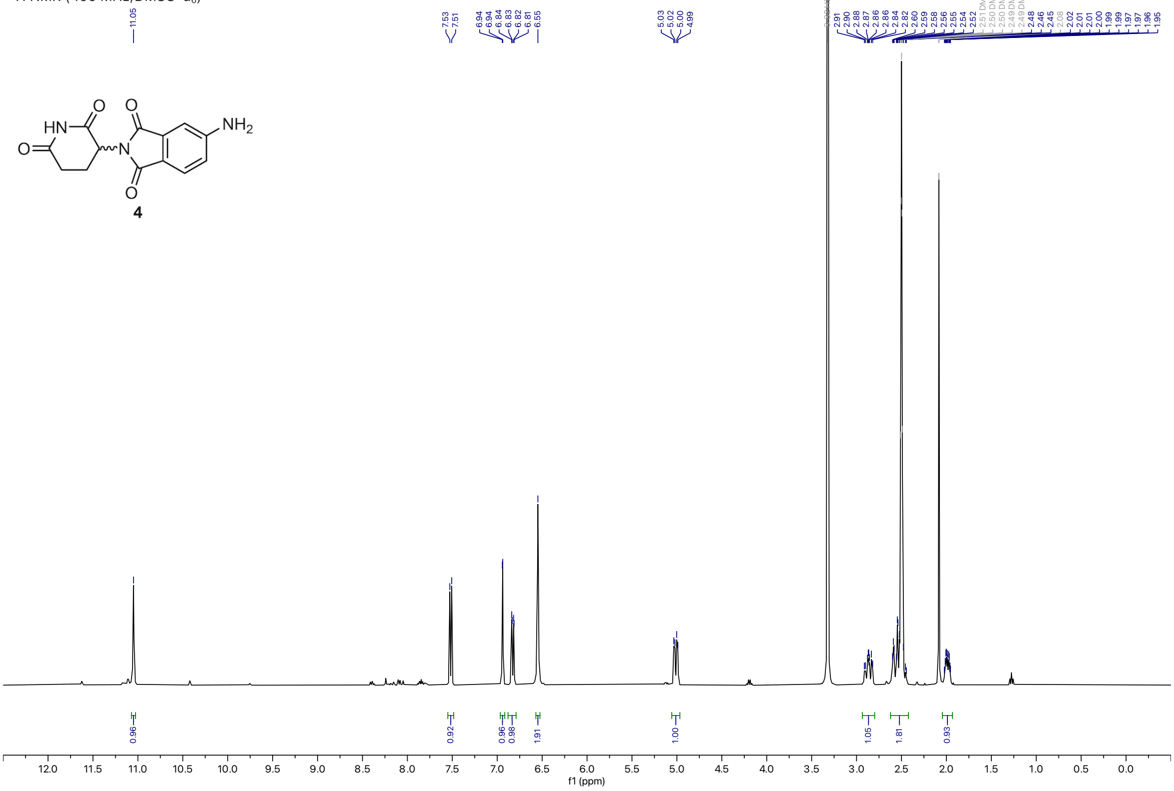


$^{13}$C NMR (101 MHz, DMSO-$d_6$)

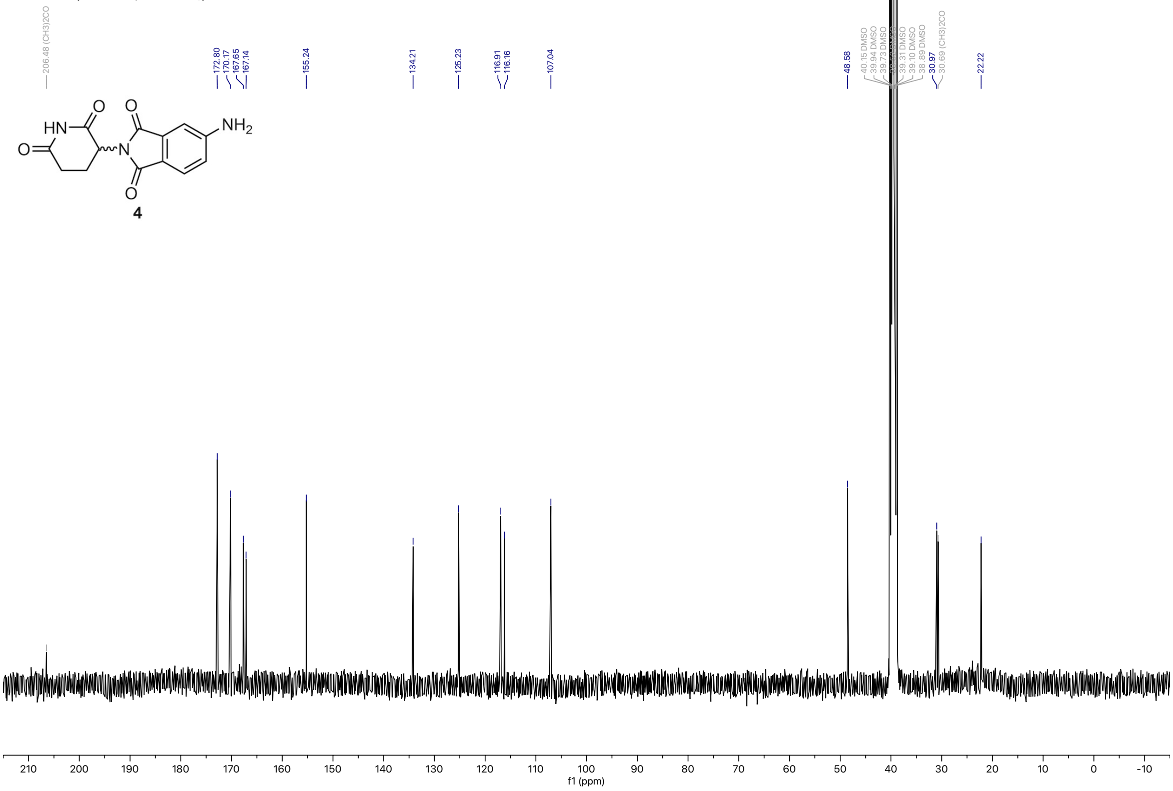

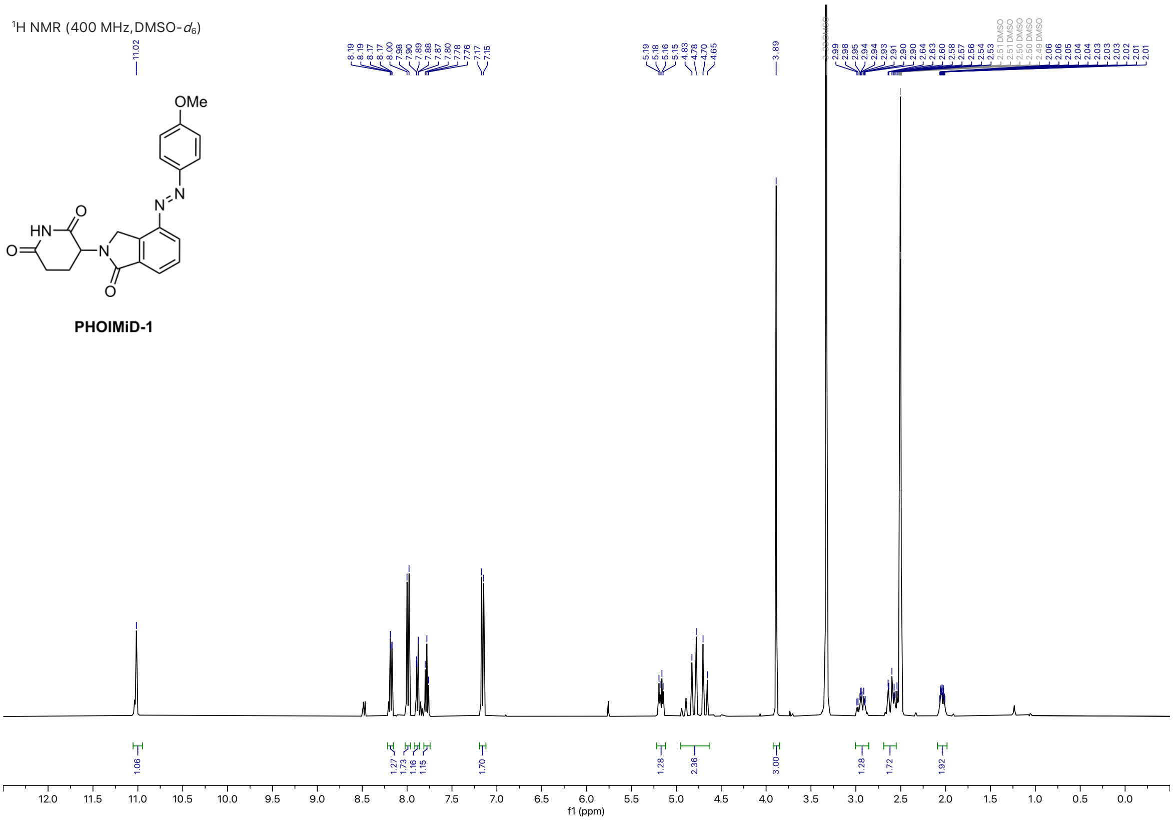
$^1$H NMR (400 MHz, DMSO-$d_6$)
OMe
PHOIMiD-1
f1 (ppm)

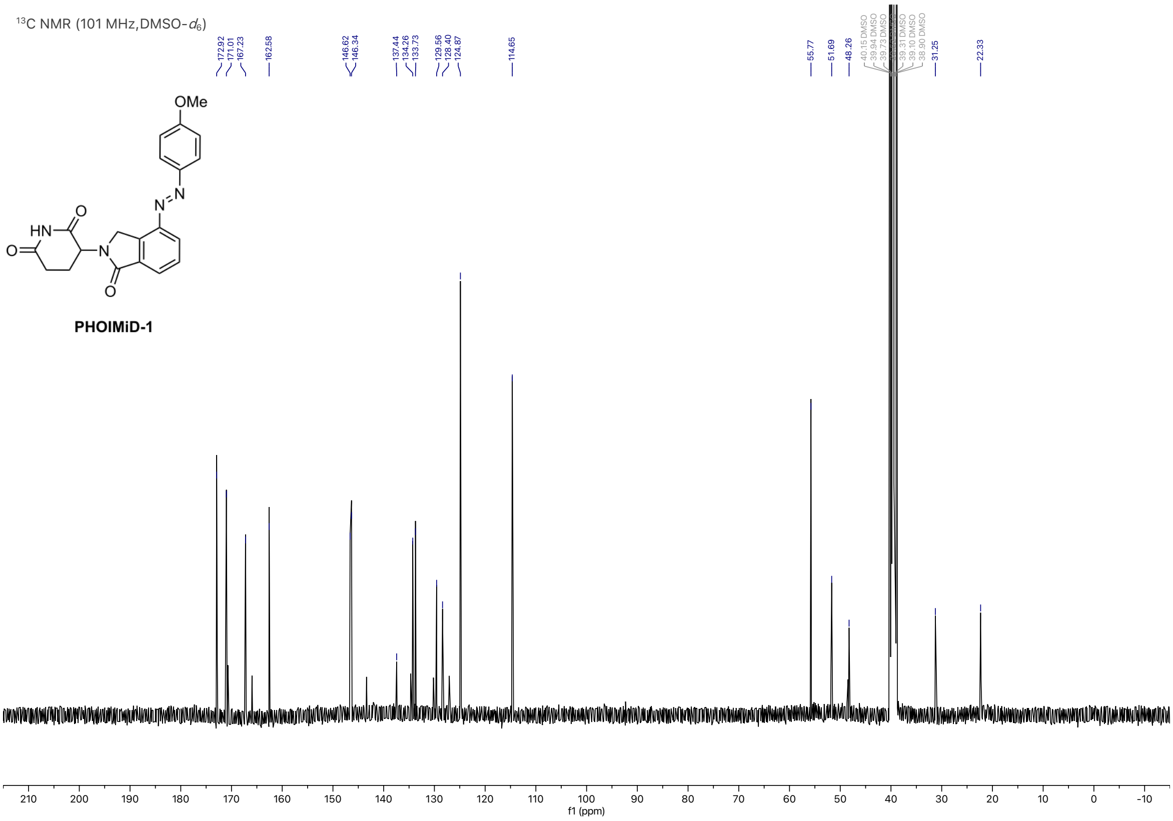
$^{13}$C NMR (101 MHz, DMSO-$d_6$)
OMe
PHOIMiD-1
f1 (ppm)

$^1$H NMR (400 MHz, DMSO-$d_6$)

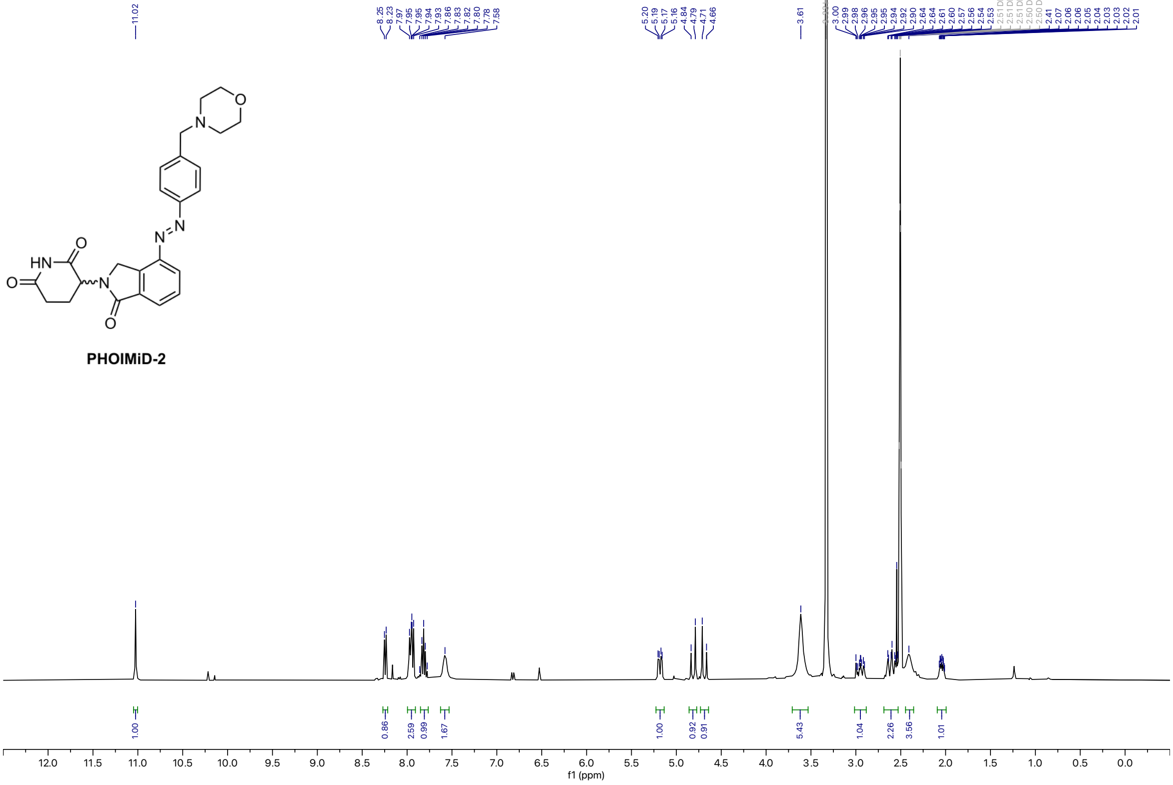


$^{13}$C NMR (101 MHz, DMSO-$d_6$)

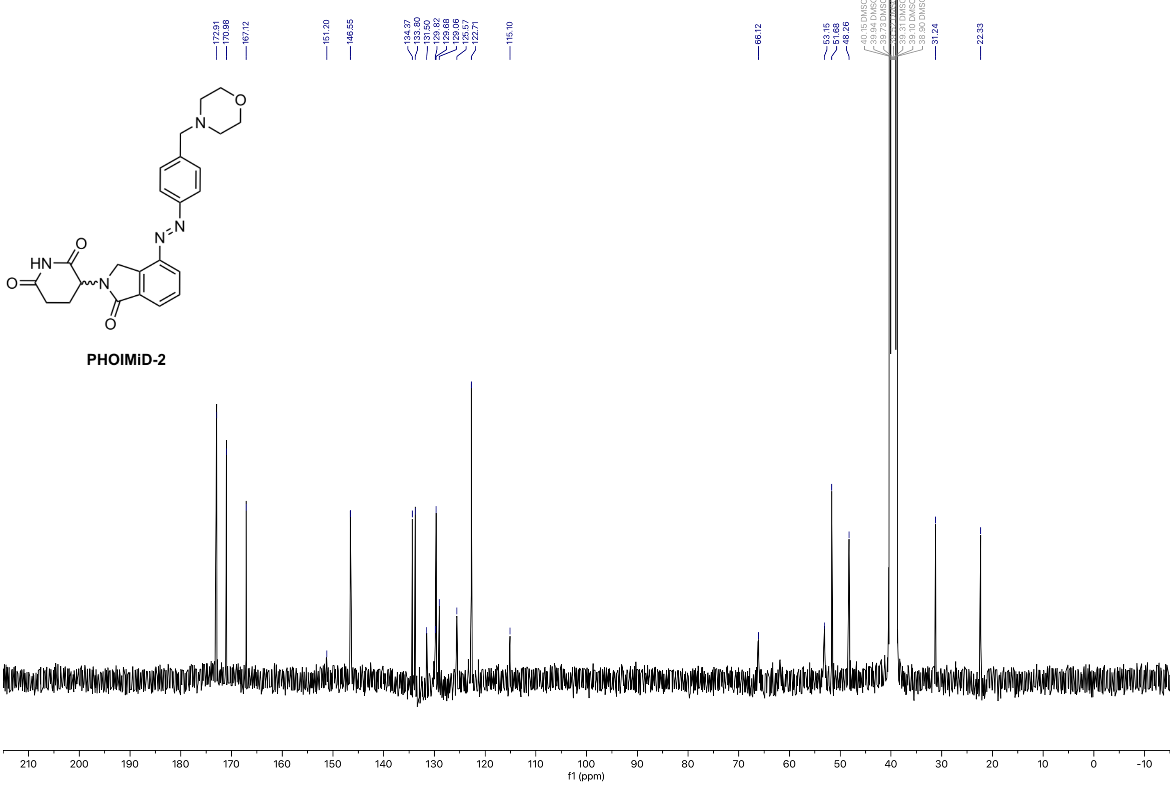

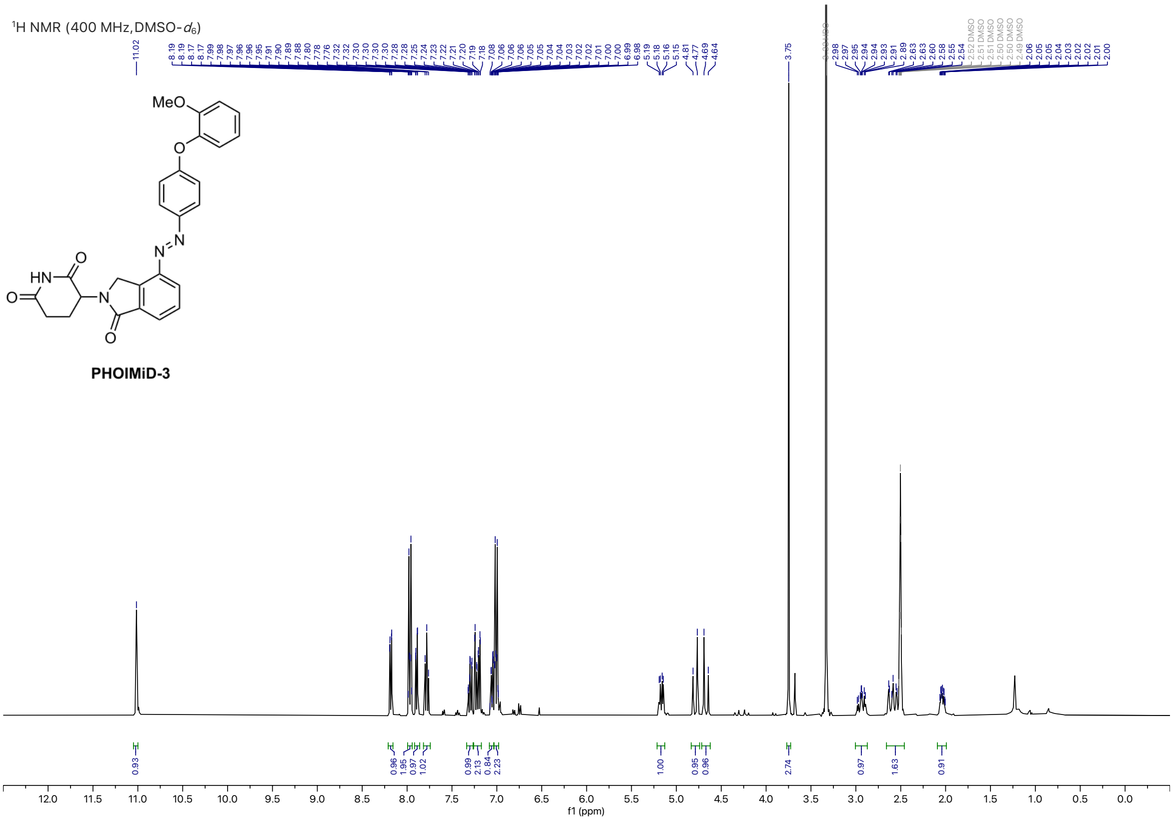


**PHOIMiD-3**

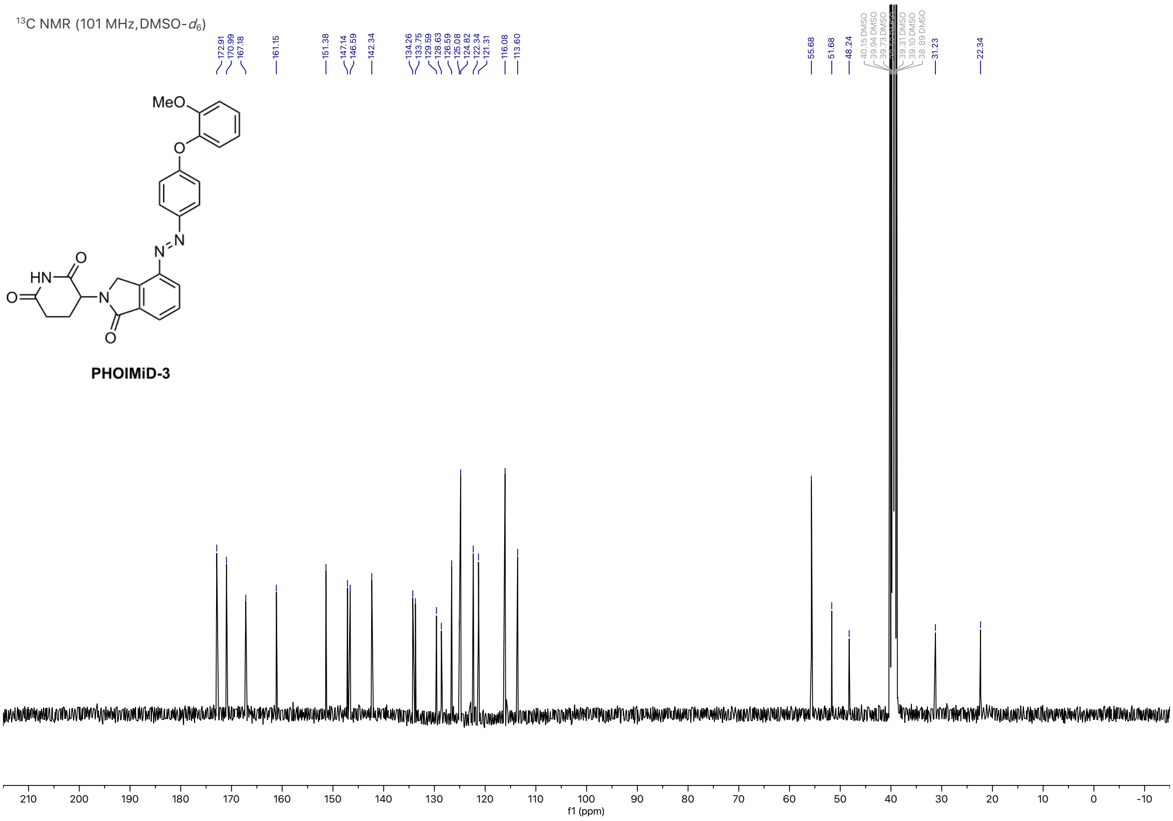


**PHOIMiD-3**

SUPPORTING INFORMATION

$^1$H NMR (400 MHz, DMSO-$d_6$)

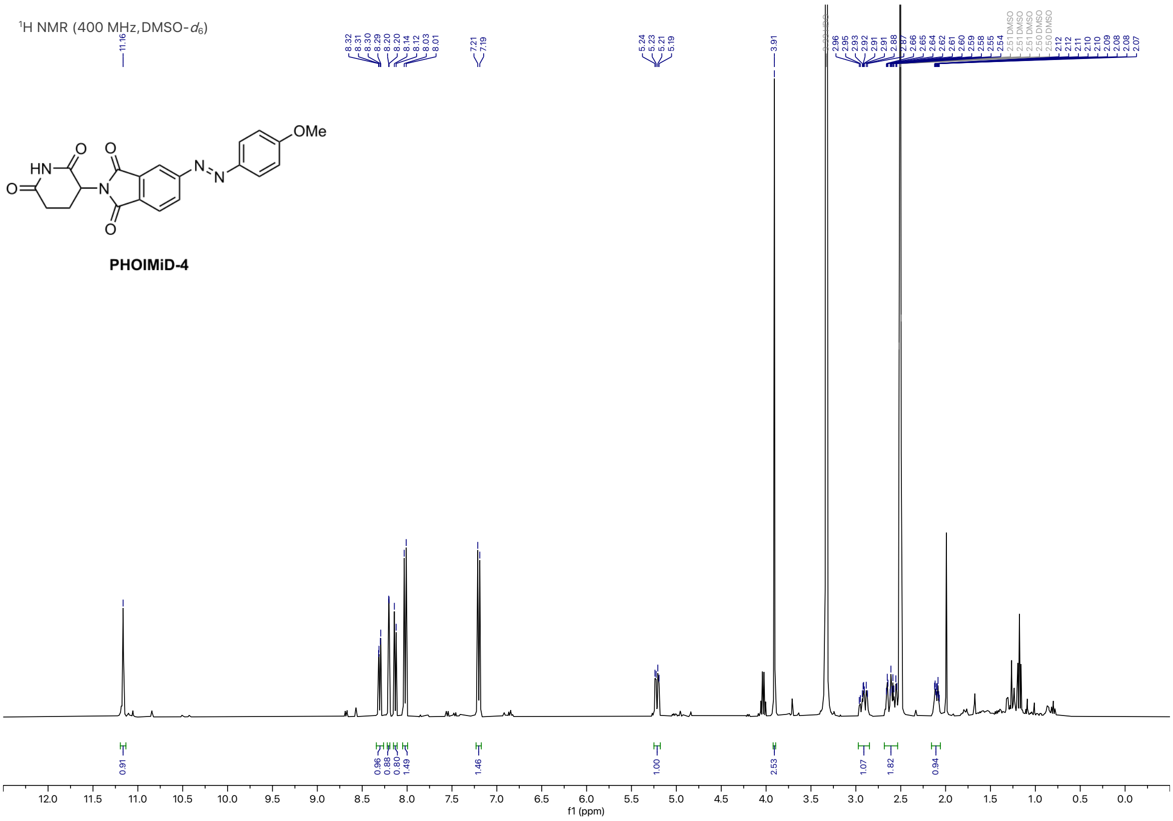


$^{13}$C NMR (101 MHz, DMSO-$d_6$)

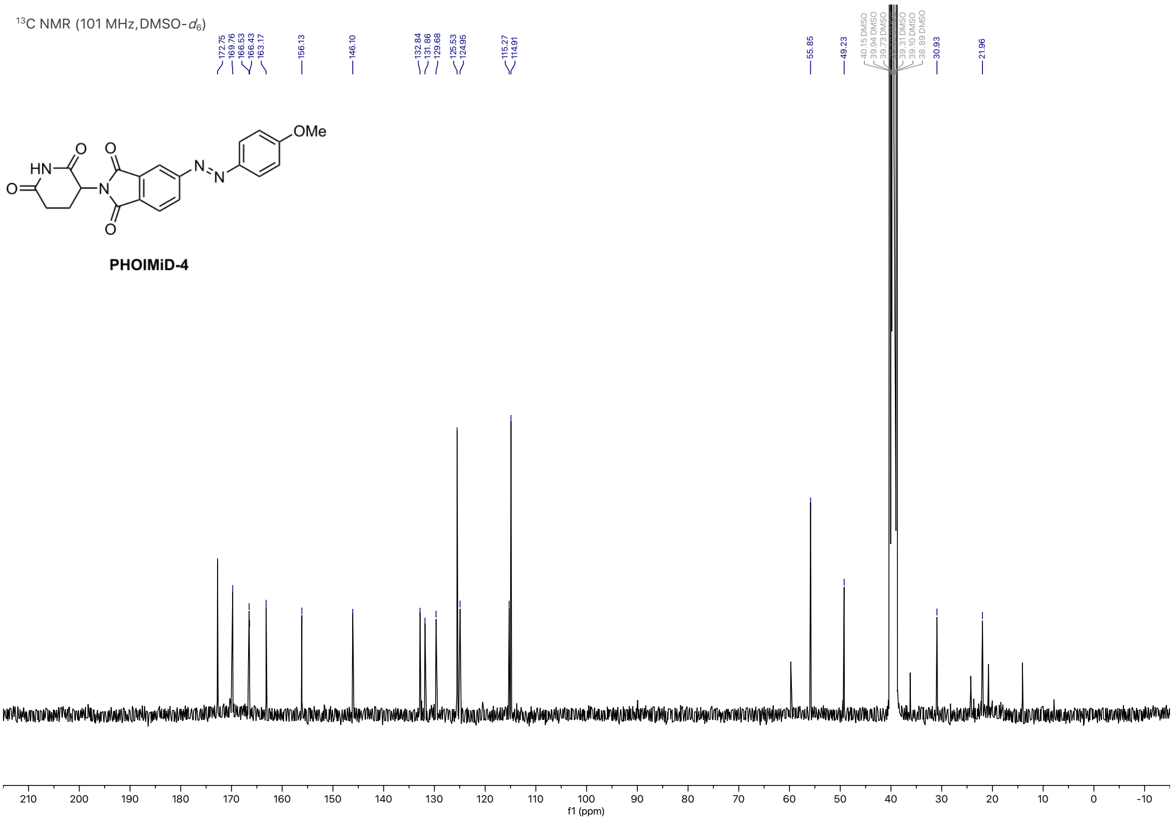

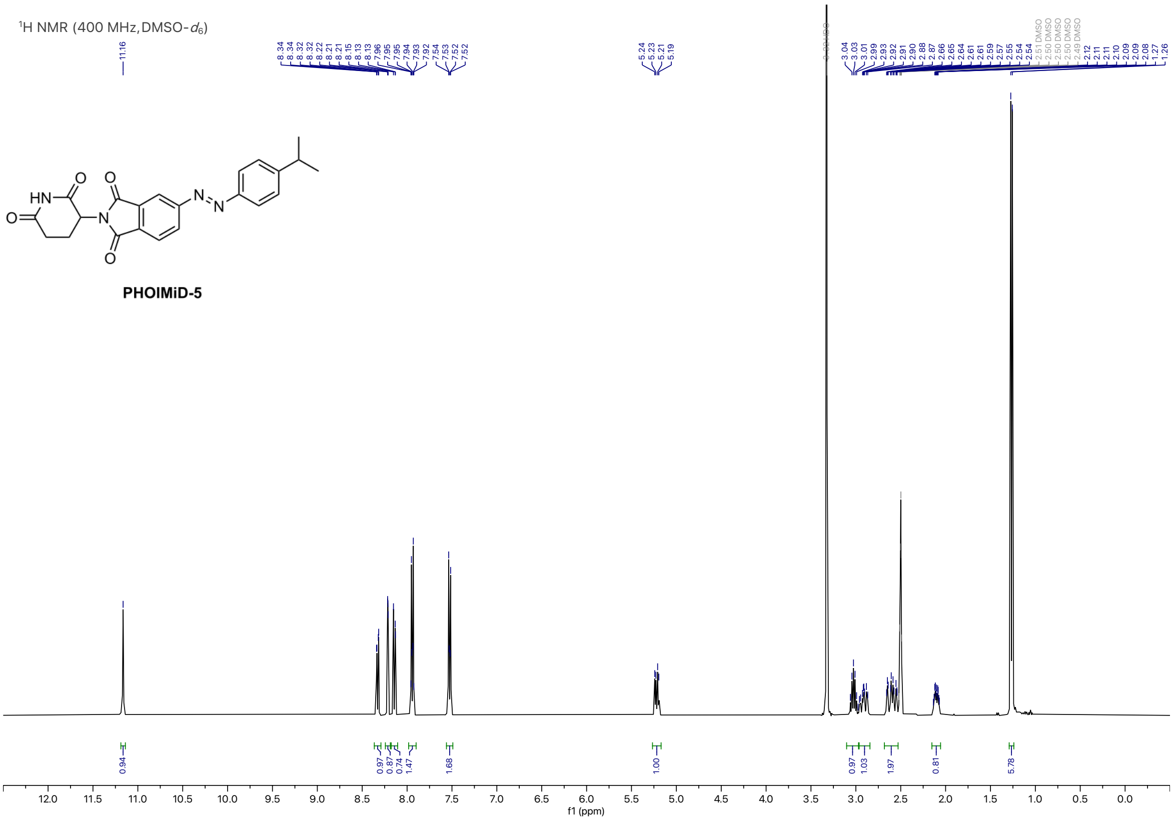
$^{1}$H NMR (400 MHz, DMSO-$d_6$)
PHOIMiD-5
f1 (ppm)

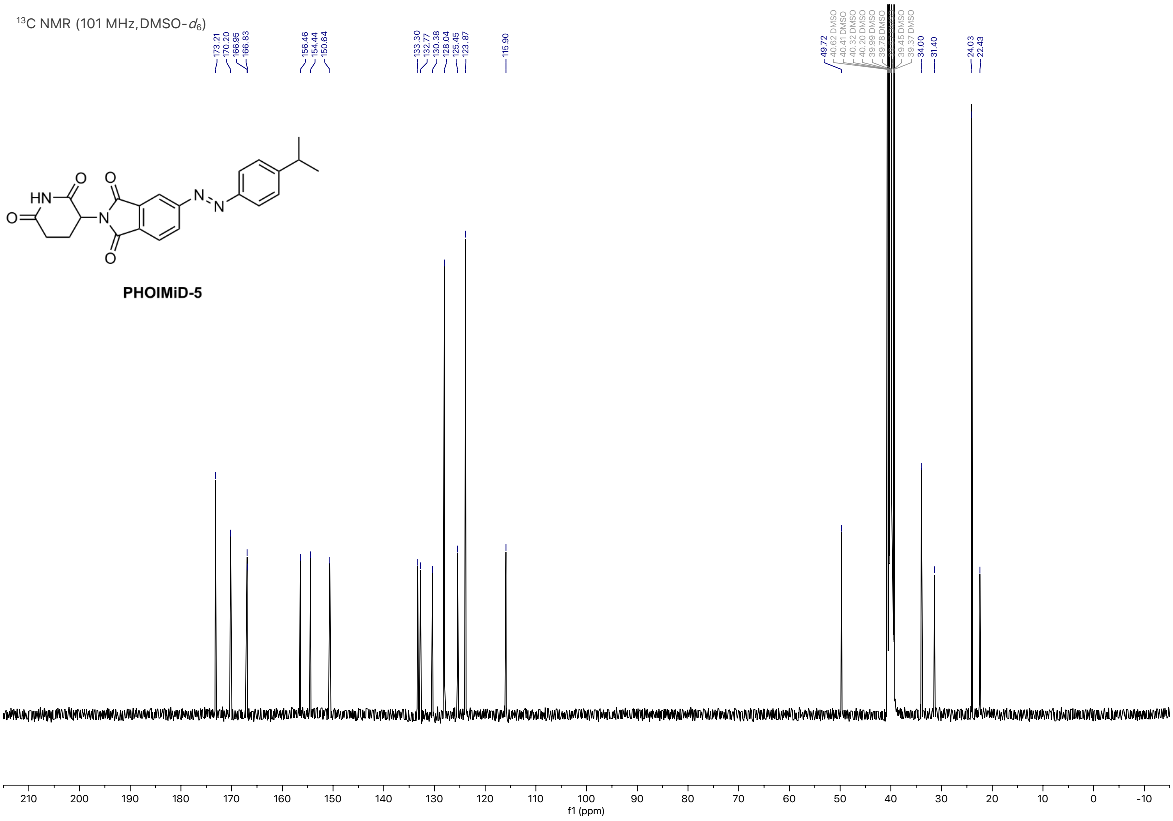
$^{13}$C NMR (101 MHz, DMSO-$d_6$)
PHOIMiD-5
f1 (ppm)

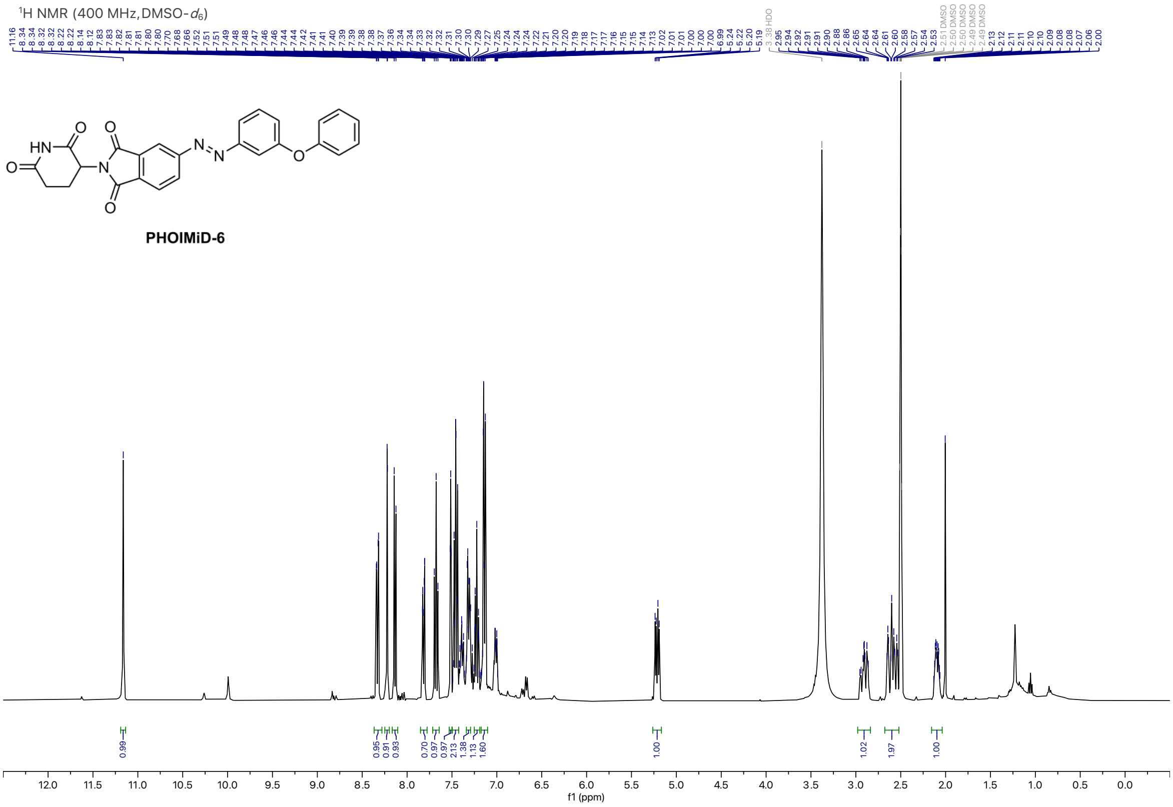

1H NMR (400 MHz, DMSO-d6)
PHOIMiD-6
f1 (ppm)


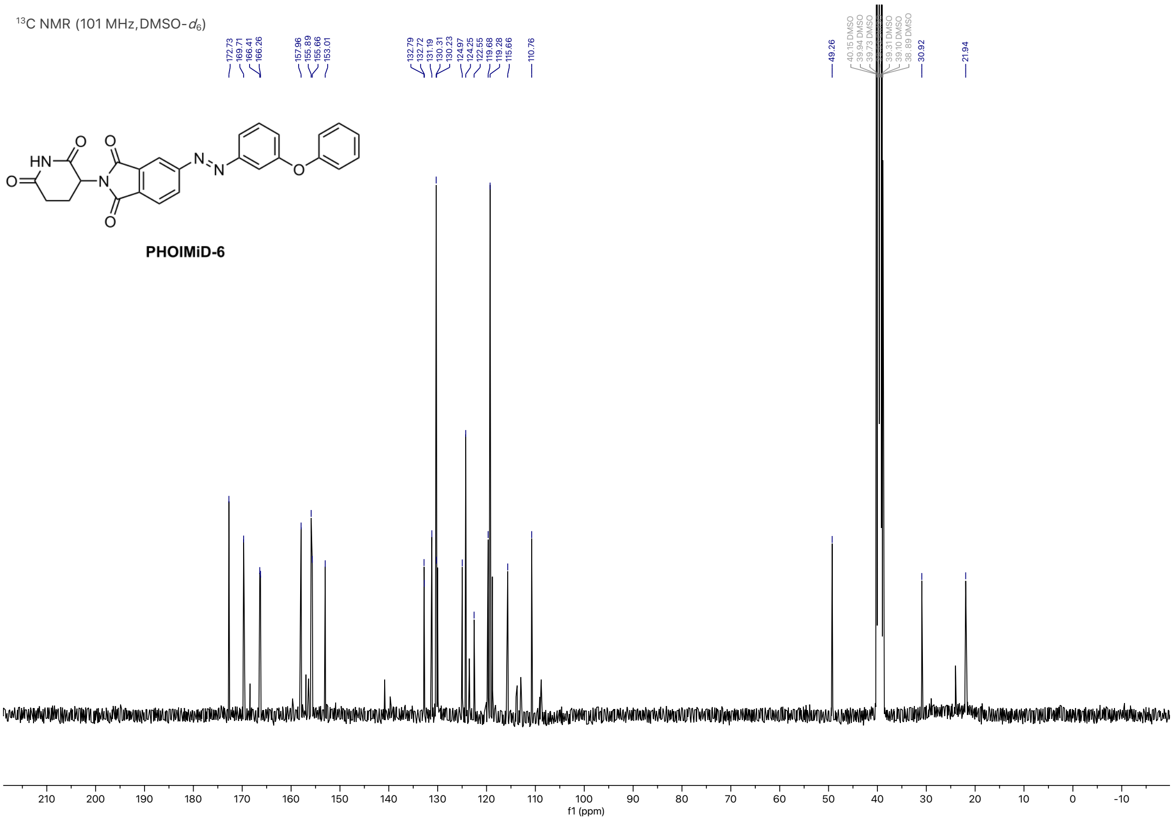

13C NMR (101 MHz, DMSO-d6)
172.73
169.71
166.41
166.26
157.96
155.89
155.66
153.01
132.79
132.72
131.19
130.31
130.23
124.97
124.25
122.55
119.68
119.28
115.66
110.76
49.26
30.92
21.94
PHOIMiD-6
f1 (ppm)

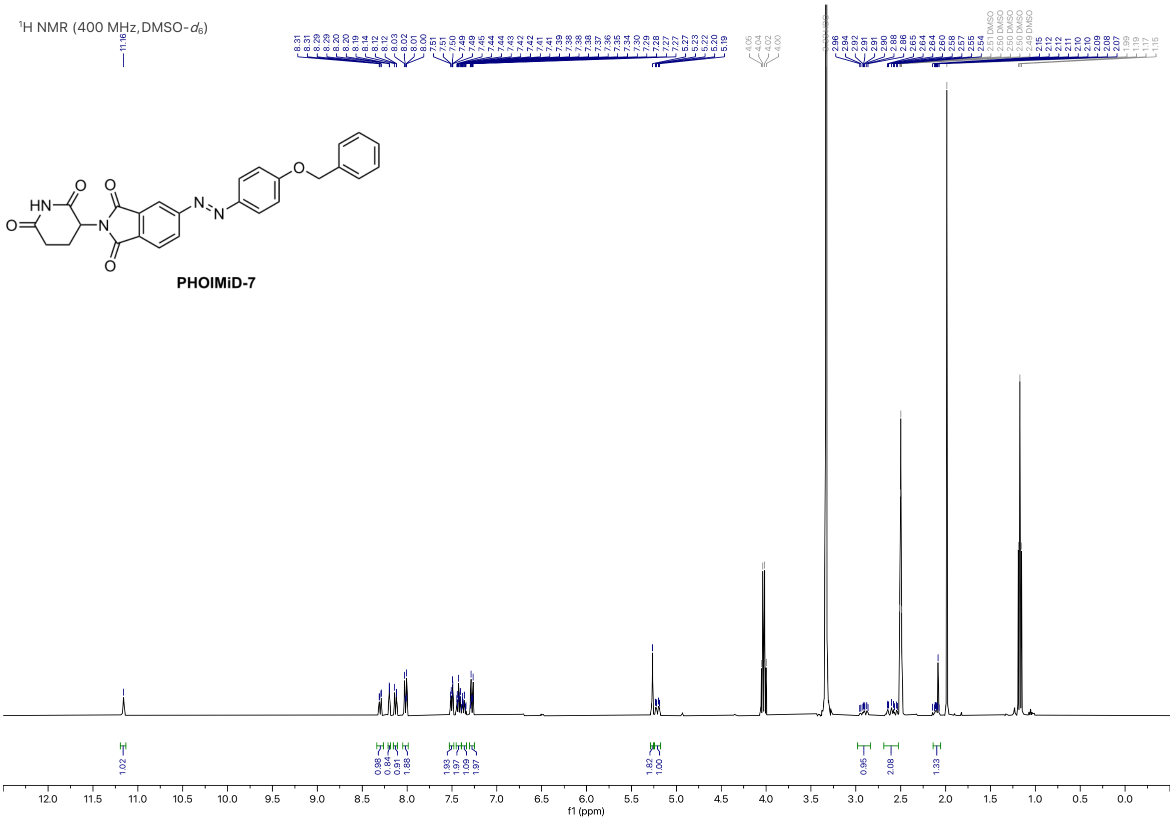
1H NMR (400 MHz, DMSO-d6)
HN
O
N
N
N
O
PHOIMiD-7
f1 (ppm)

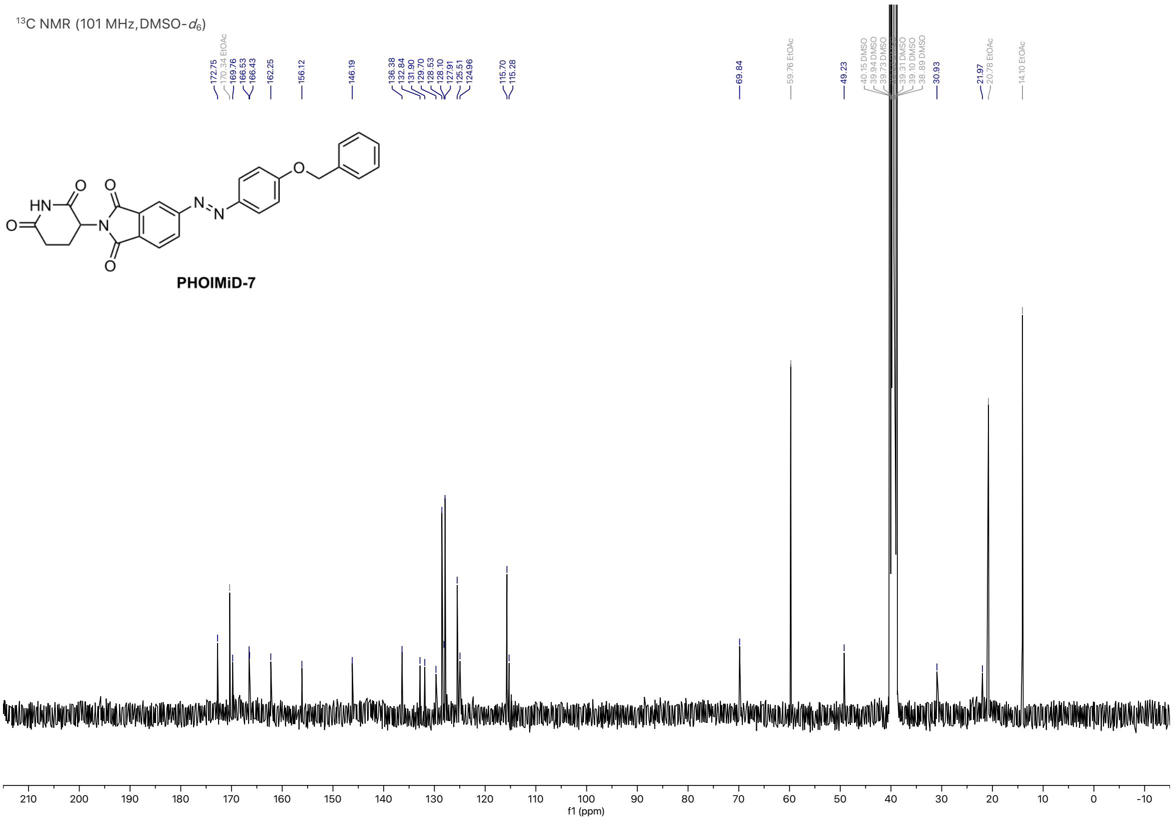
13C NMR (101 MHz, DMSO-d6)
172.75
170.34 EtOAc
169.76
166.53
166.43
162.25
156.12
146.19
136.38
132.84
131.90
129.70
128.53
128.10
127.91
125.51
124.96
115.70
115.28
69.84
59.76 EtOAc
49.23
30.93
21.97
20.78 EtOAc
14.10 EtOAc
PHOIMiD-7
f1 (ppm)

$^{1}H$ NMR (400 MHz, DMSO-$d_6$)

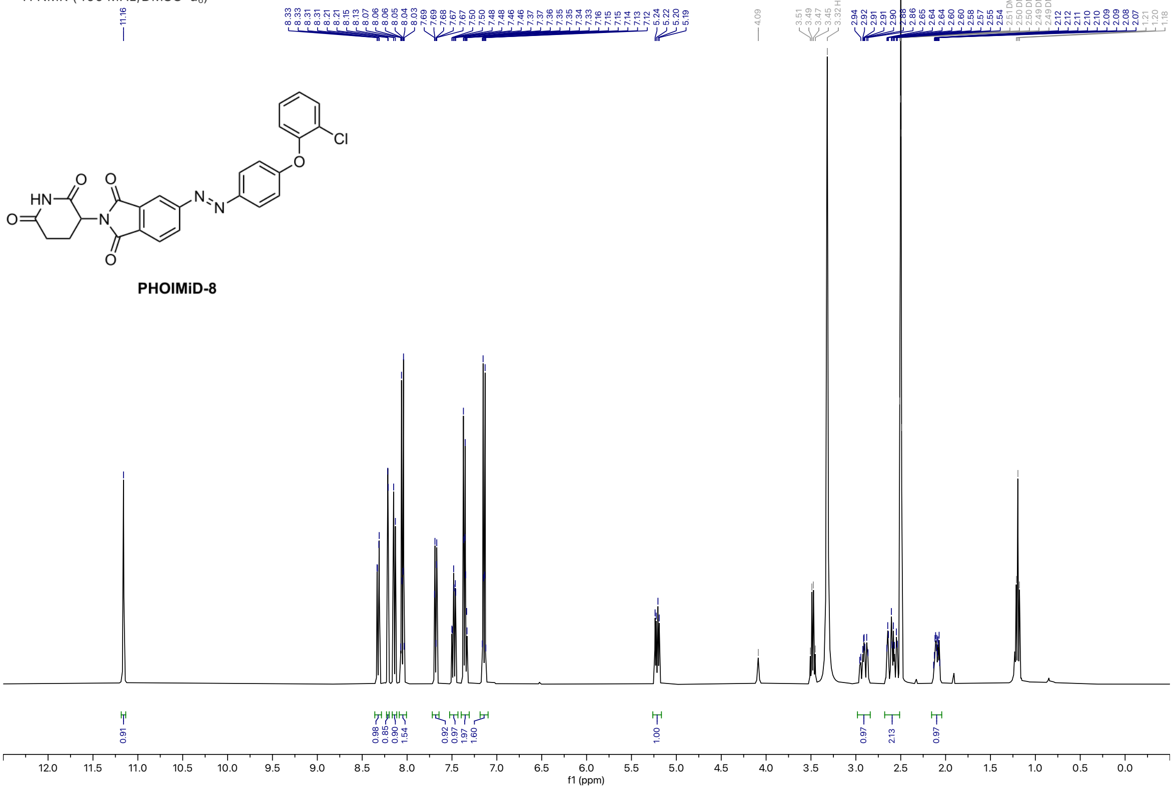


$^{13}C$ NMR (101 MHz, DMSO-$d_6$)

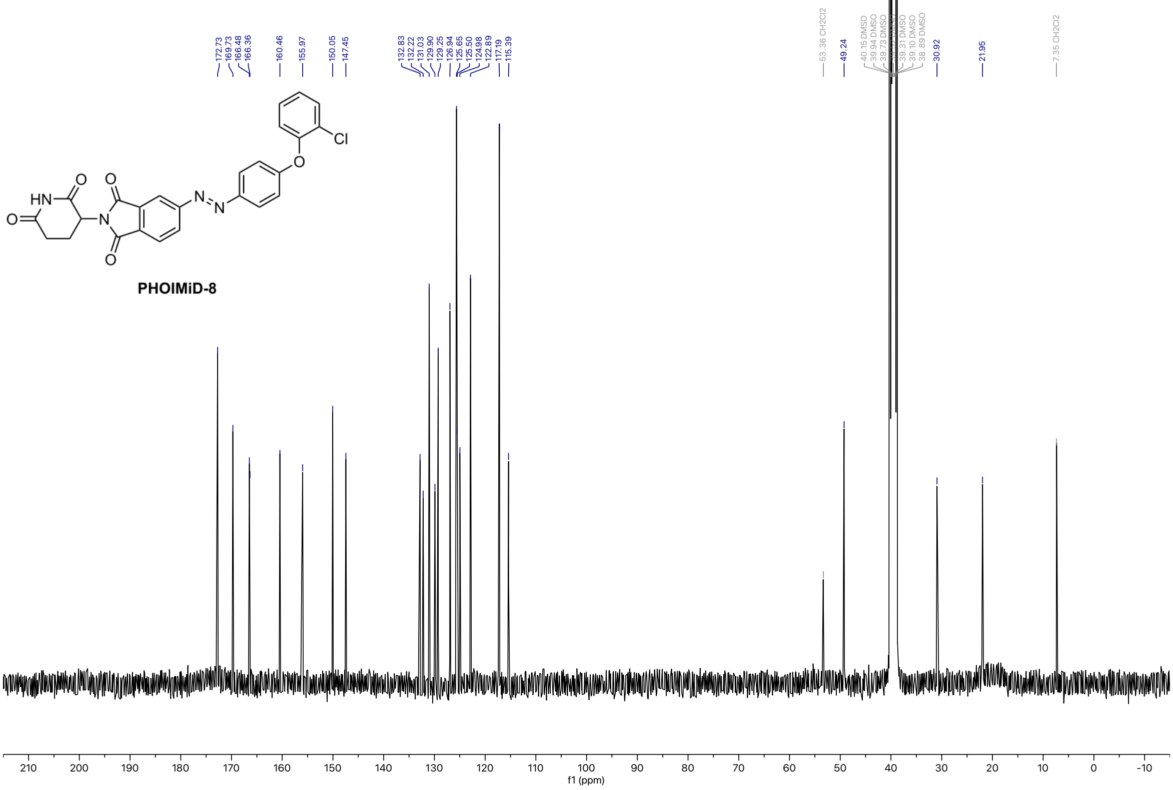

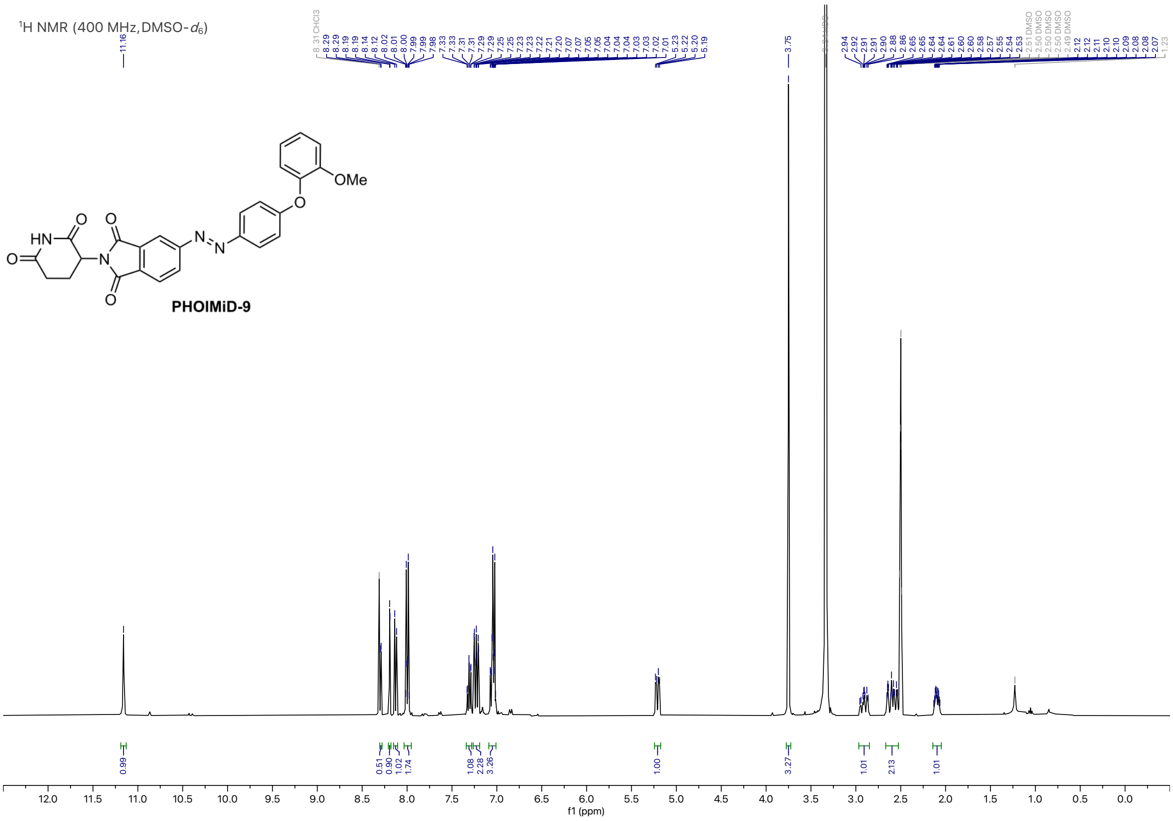

$^{1}$H NMR (400 MHz, DMSO-$d_6$)
OMe
HN
O
N
N
N
PHOIMiD-9
f1 (ppm)


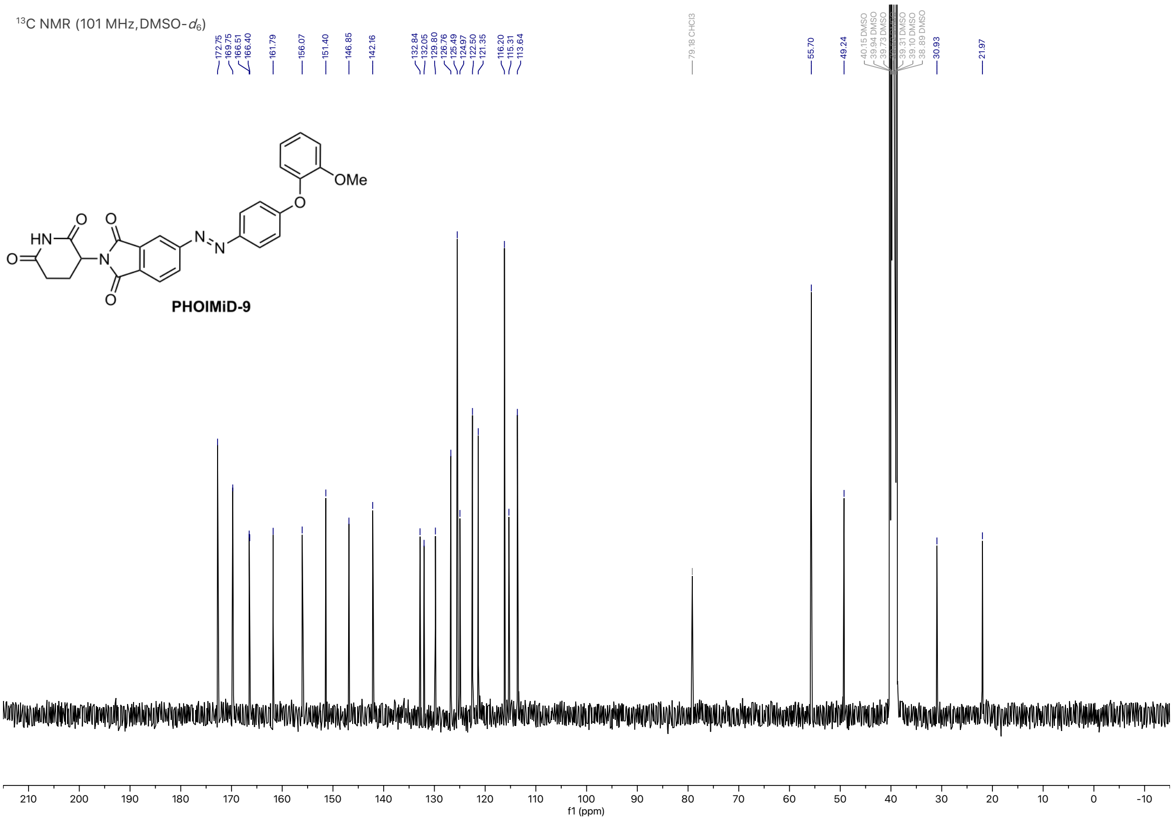

$^{13}$C NMR (101 MHz, DMSO-$d_6$)
172.75
169.75
166.51
166.40
161.79
156.07
151.40
146.85
142.16
132.84
132.05
129.80
126.76
125.49
124.97
122.50
121.35
116.20
115.31
113.64
79.18 CHCl3
55.70
49.24
30.93
21.97
OMe
PHOIMiD-9
f1 (ppm)

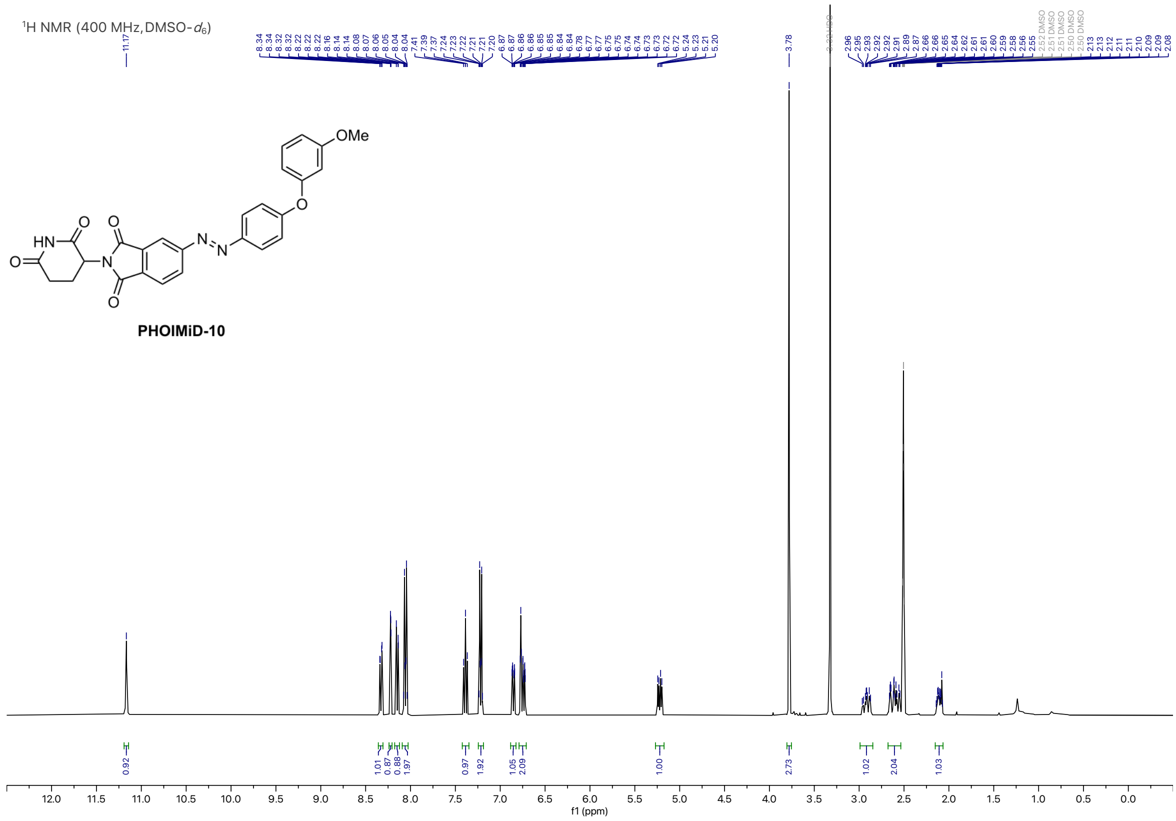

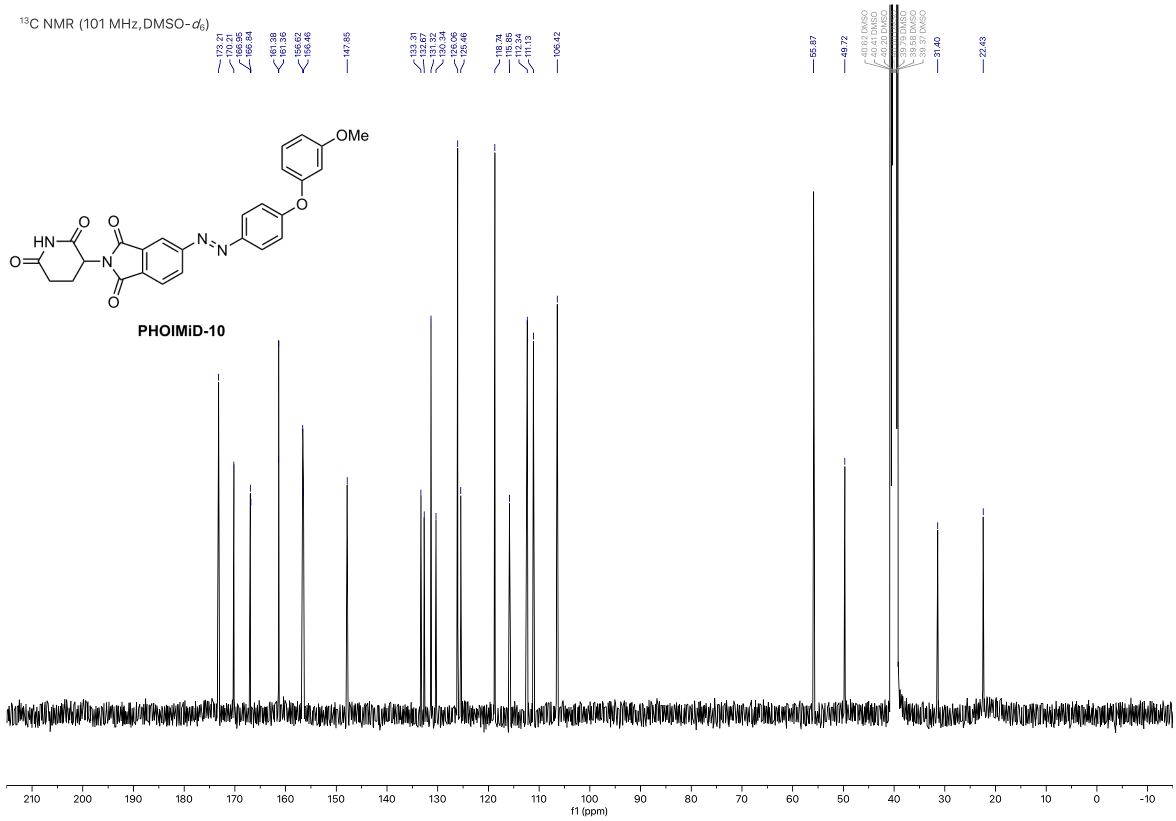

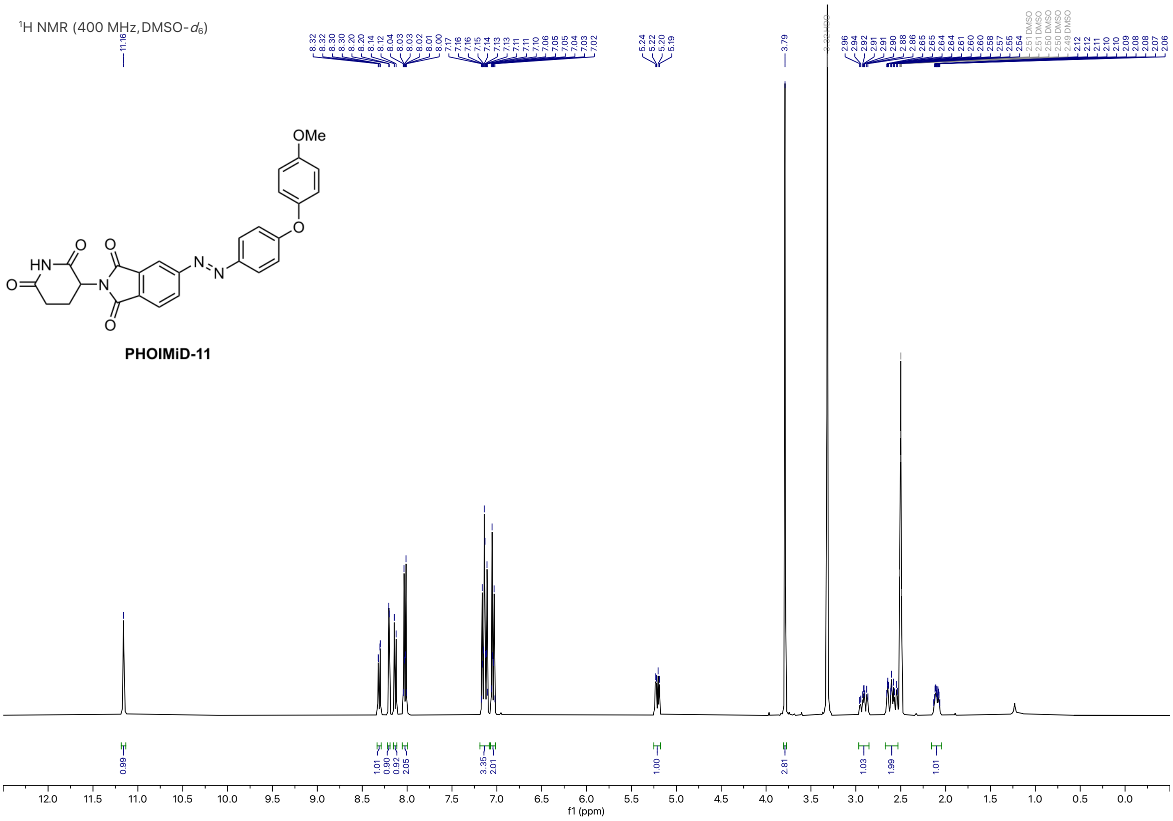
¹H NMR (400 MHz, DMSO-d6)
PHOIMiD-11
f1 (ppm)

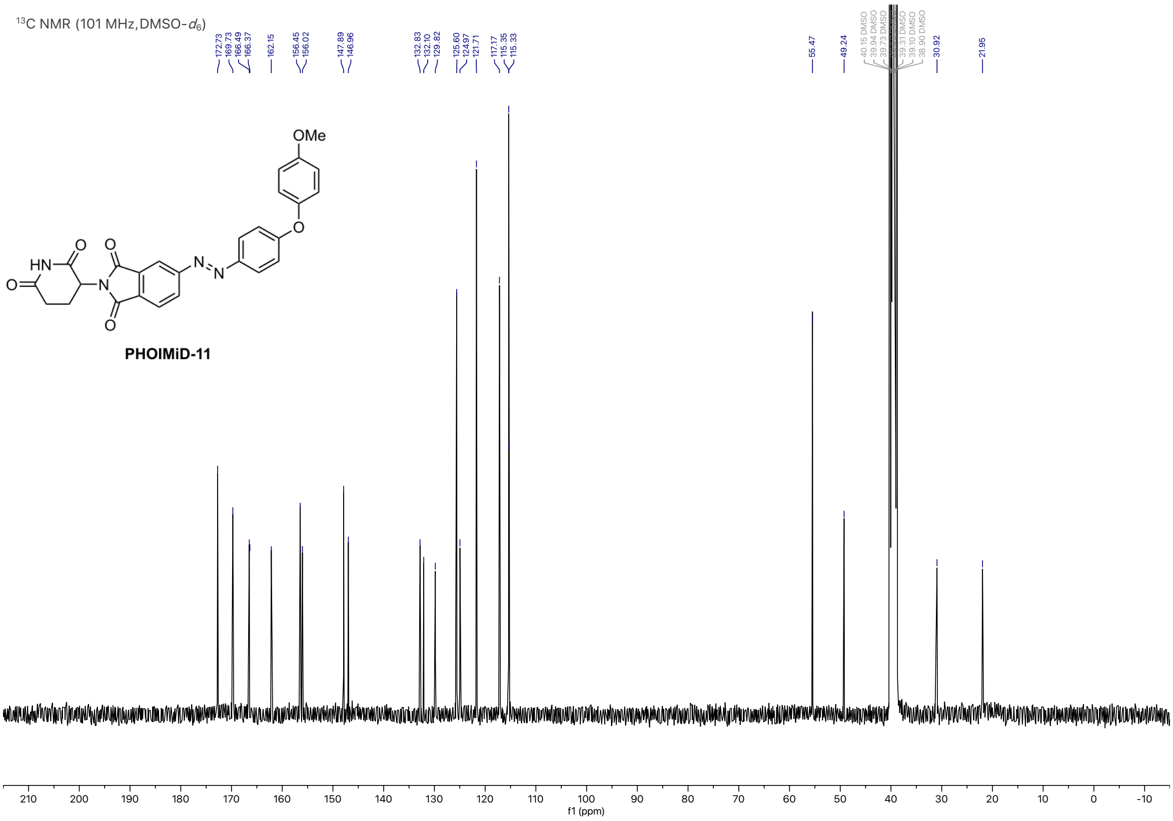
¹³C NMR (101 MHz, DMSO-d6)
PHOIMiD-11
f1 (ppm)
